# Racial differences in androgen metabolism and receptor signaling in prostate cancer

**DOI:** 10.1101/2021.03.31.437727

**Authors:** Swathi Ramakrishnan, Eduardo Cortes-Gomez, Kristopher Attwood, Rick A Kittles, Jianmin Wang, Spencer R Rosario, Dominic J Smiraglia, Gissou Azabdaftari, James L Mohler, Wendy J Huss, Anna Woloszynska

## Abstract

Dihydrotestosterone (DHT) and testosterone (T) mediated androgen receptor (AR) nuclear translocation initiates transcription of AR target genes that are pivotal for prostate cancer (PrCa) development and progression. Here we provide data indicating that in contrast to European American (EA) men, African American (AA) men with localized PrCa can exploit an alternative progesterone-androsterone-5α-androstanedione pathway for DHT biosynthesis. Enzymes that are involved in alternate pathways of DHT biosynthesis are elevated in PrCa tissues from AA men, compared to EA men, and also correlated with increased serum DHT levels. In addition, higher serum DHT levels reflect increased RNA expression of AR target genes in PrCa tissues from AA men. Interestingly, serum T but not DHT levels are significantly lower in AA men compared to EA men with PrCa. Furthermore, serum progesterone and related intermediate metabolites levels that are produced during alternate pathways of DHT biosynthesis are significantly lower in AA men with PrCa and associated with a shorter time to disease progression. These data highlight that androgen biosynthesis is altered in therapy naïve localized PrCa in AA men, and can potentially serve as prognostic indicators of disease progression.

**Significance:** Our work provides a rationale to examine potential pharmacological interventions that target androgen biosynthesis and AR signaling earlier in the disease continuum in AA men with PrCa. Additionally, our study lays the groundwork for developing serum measurements of intermediate androgen metabolites as PrCa prognostic biomarkers.

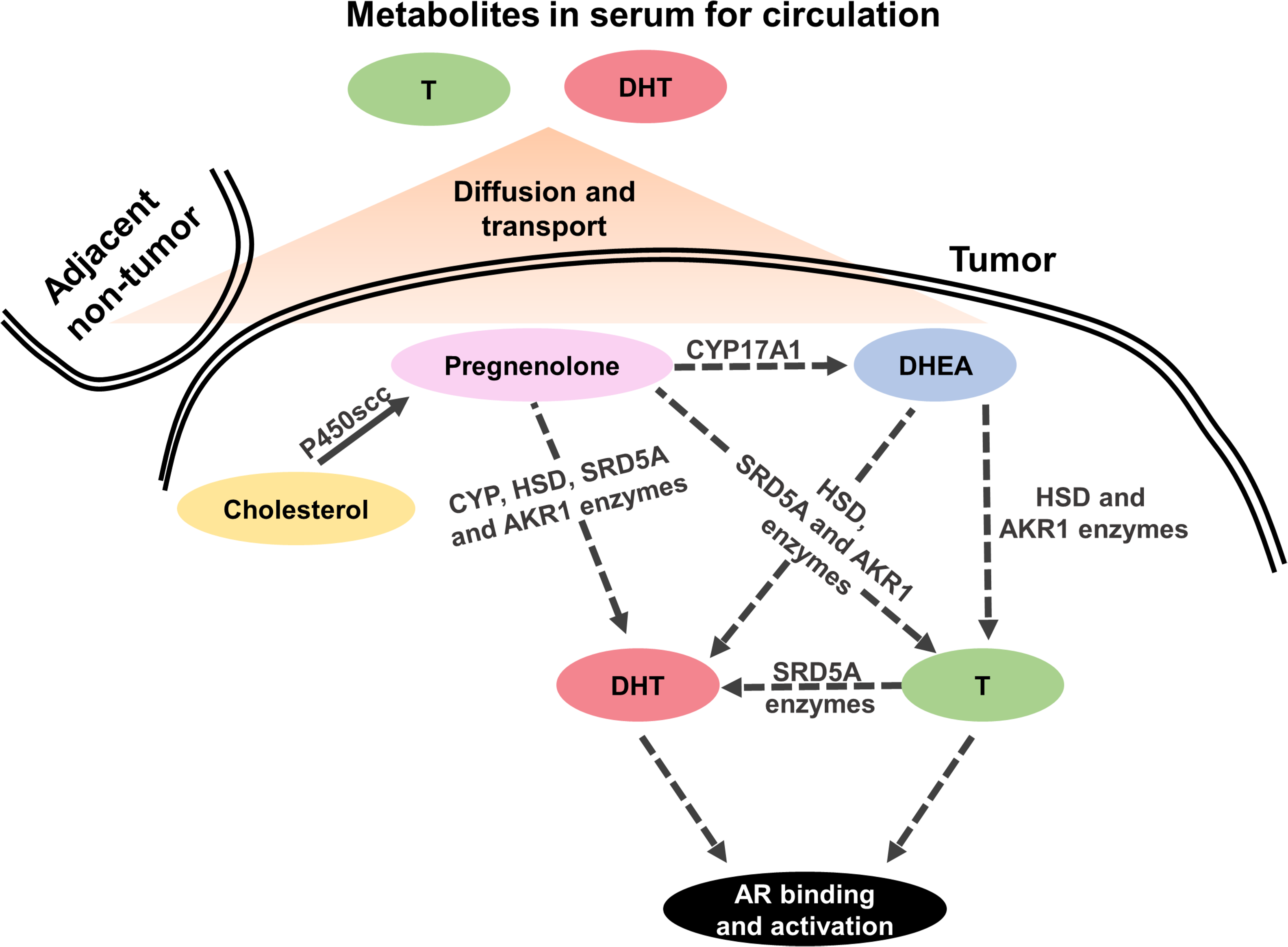

## Introduction

Hormone-driven androgen receptor (AR) and its transcriptional activity is a focal signaling axis in prostate cancer (PrCa)^1–3^. Androgens, including dihydrotestosterone (DHT) and testosterone (T), are synthesized through the conversion of different intermediate metabolites that are regulated by several enzymes ^4^. DHT and T bind to and initiate AR nuclear translocation resulting in the transcription of several AR target genes that control the cell cycle, apoptosis, differentiation, and other processes necessary for PrCa development and progression^2, 5–11^. Therefore, both androgen synthesis and AR activity are the primary targets of systemic therapies used in PrCa^1–3^. PrCa from African American (AA) men have increased AR transcriptional activity compared to European American (EA) men. Importantly, the association between DHT or T levels in the serum and AR expression/activity in PrCa tissue from AA men has not been established^12, 13^. In EA men, there is some evidence of a negative correlation between T levels in the serum and AR protein expression but not transcriptional activity in PrCa tissues^14^.

DHT and T are primarily produced in the Leydig cells of the testis, with small quantities being synthesized in the adrenal glands^15, 16^. In the non-diseased state, AA men have increased DHT levels in the serum compared to EA men^3, 17–20^. Prostate tumors themselves are also known to produce their own androgens, especially in the setting of androgen deprivation therapies^21–26^. Irrespective of the location, DHT and T *de novo* synthesis requires cholesterol as the primary precursor^4^. In AA men, high levels of total cholesterol in the serum are associated with high-risk PrCa disease and are considered a significant risk factor for PrCa recurrence^27, 28^. Whether serum cholesterol levels are indicative of increased DHT and T biosynthesis is unknown. Cholesterol is converted to pregnenolone that can lead to T synthesis through three possible pathways: (1) dehydroepiandrosterone (DHEA)-androstenedione (ASD), (2) DHEA-androstenediol, or (3) progesterone-ASD conversion^1, 29, 30^. Similarly, DHT can be synthesized through three pathways that involve: (1) DHEA-T, (2) progesterone-androsterone (AND)-androstanedione (5α-dione), or (3) DHEA-ASD-5α-dione^1, 29, 30^. Compared to EA men, PrCa tissues from AA men have significantly higher levels of intermediate androgen metabolites, such as DHEA and ASD but not DHT or T^31, 32^. Whether there are race-specific differences in DHT, T, and intermediate metabolite levels in the adjacent non-tumor tissues is unknown. The enzymes regulating DHT and T synthesis include Cytochrome P450 17A1 (CYP17A11), 3β-hydroxysteroid dehydrogenase (3βHSD), and Aldo-keto reductase family 1 (AKR1C)^4^. Thus far, only a single study has reported race-specific differences in the RNA expression of enzymes involved in androgen metabolism^33^. These enzymes are also known to harbor single nucleotide polymorphisms (SNPs) found at different frequencies in AA and EA men in healthy and diseased states^24, 34–38^. Although the functional consequences of these SNPs remain poorly understood, they are associated with clinicopathological parameters and occur at different frequencies in diverse populations^24, 34–43^.

There are limited comparative studies of androgens and intermediate metabolites in the serum of AA and EA men at initial PrCa diagnosis and before radical prostatectomy (RP)^18, 44–46^. Here, we investigate differences in levels of androgen metabolites in the serum in relation to clinical outcomes and AR expression/activity of AA and EA men at the time of PrCa diagnosis. Our results show that the expression of enzymes that synthesize DHT through alternative pathways is increased in PrCa from AA men compared to EA men. We show for the first time that the expression of enzymes involved in the alternative DHT synthesis pathways correlates with DHT levels in the serum of AA but not EA men with PrCa. We also find that DHT levels in the serum correlate to AR transcriptional activity in PrCa tissues from AA but not EA men. Interestingly, in the serum, low levels of T and a subset of intermediate metabolites more commonly found in AA men with PrCa associate with shorter time to disease progression. Finally, we identified a novel SNP in the CYP11B family of enzymes that lead to aldosterone synthesis at an increased frequency in AA men with PrCa. In summary, our data indicate that AA men more frequently utilize alternative pathways of DHT synthesis, and that the levels of androgen intermediate metabolites in the serum at the time of RP can be indicators of clinical outcomes in AA and EA men.

## Results

### Androgen synthesis enzymes from the alternate pathway are upregulated in PrCa from AA men

To investigate racial disparities at the time of RP, we utilized an unbiased RNA sequencing approach to determine differences in the cellular transcriptome in PrCa and adjacent non-tumor tissues from AA men and EA men. We measured differences in clinical samples hereafter referred to as Roswell Park Cohort 1 (**Supplementary Table 1**). RNA sequencing analyses revealed that close to 50% of the top 500 upregulated and 500 downregulated genes are protein coding genes when comparing PrCa from AA and EA men in Roswell Park Cohort 1 (**Figure 1A**, Demographics in **Supplementary Table 2**). Interestingly, the remainder of the dysregulated genes belong to families of poorly understood long non-coding RNAs, processed pseudogenes, and antisense RNAs (**Figure 1A**). We investigated pathways that are enriched in PrCa derived from AA men compared to EA men using Genego MetaCore. These analyses revealed that androgen metabolism is one of the top 5 pathways upregulated in PrCa derived from AA men compared to EA men Roswell Park Cohort 1 (**Figure 1B, Supplementary Table 3**). Specifically, the AKR1C1 that produces DHT using alternative pathways were upregulated in PrCa and adjacent non-tumor tissue AA men compared to EA men (**Figure 2A, Supplementary Table 3**). We validated the expression of a subset of these enzymes in matched clinical samples from AA (n=5) and EA men (n=4) (**Figure 2B-C**, **Supplementary Figure 1**, primers listed in **Supplementary Table 4**).

**Figure 1.**
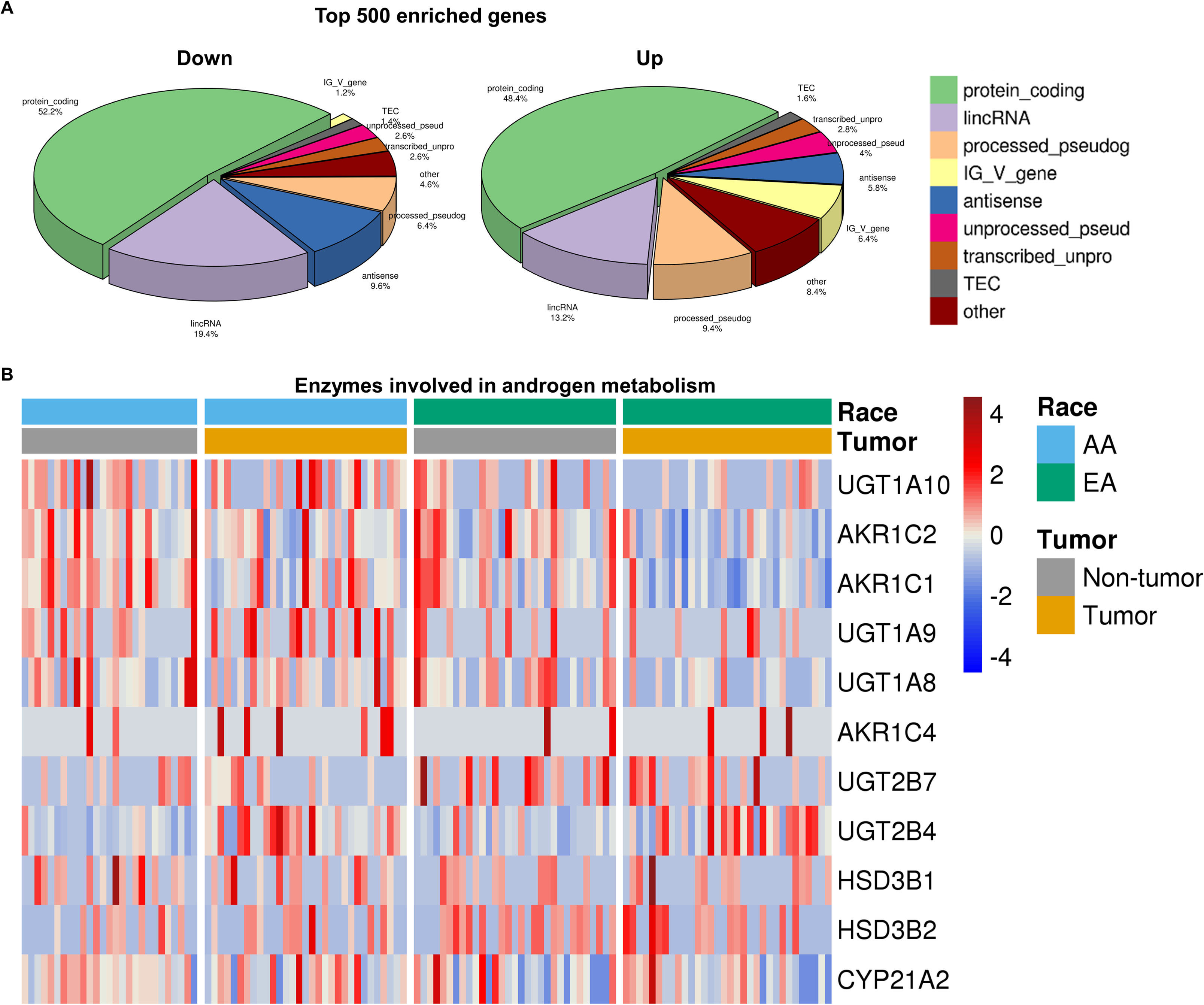
Enzymes involved in DHT biosynthesis from alternate pathways are overexpressed in PrCa from AA men. **A.** Pie charts represent the distribution of the type of gene categories that are differentially expressed in PrCa tissues from AA men compared to EA men from Roswell Park Cohort 1. The left pie chart represents the top 500 downregulated genes in PrCa tissues from AA men. The right pie chart represents the top 500 upregulated genes in PrCa tissues from AA men**. B.** The heatmap represents a subset of enzymes involved in alternate pathways of androgen biosynthesis that are differentially expressed in PrCa and adjacent non-tumor tissues from AA and EA men in Roswell Park Cohort 1.

**Figure 2.**
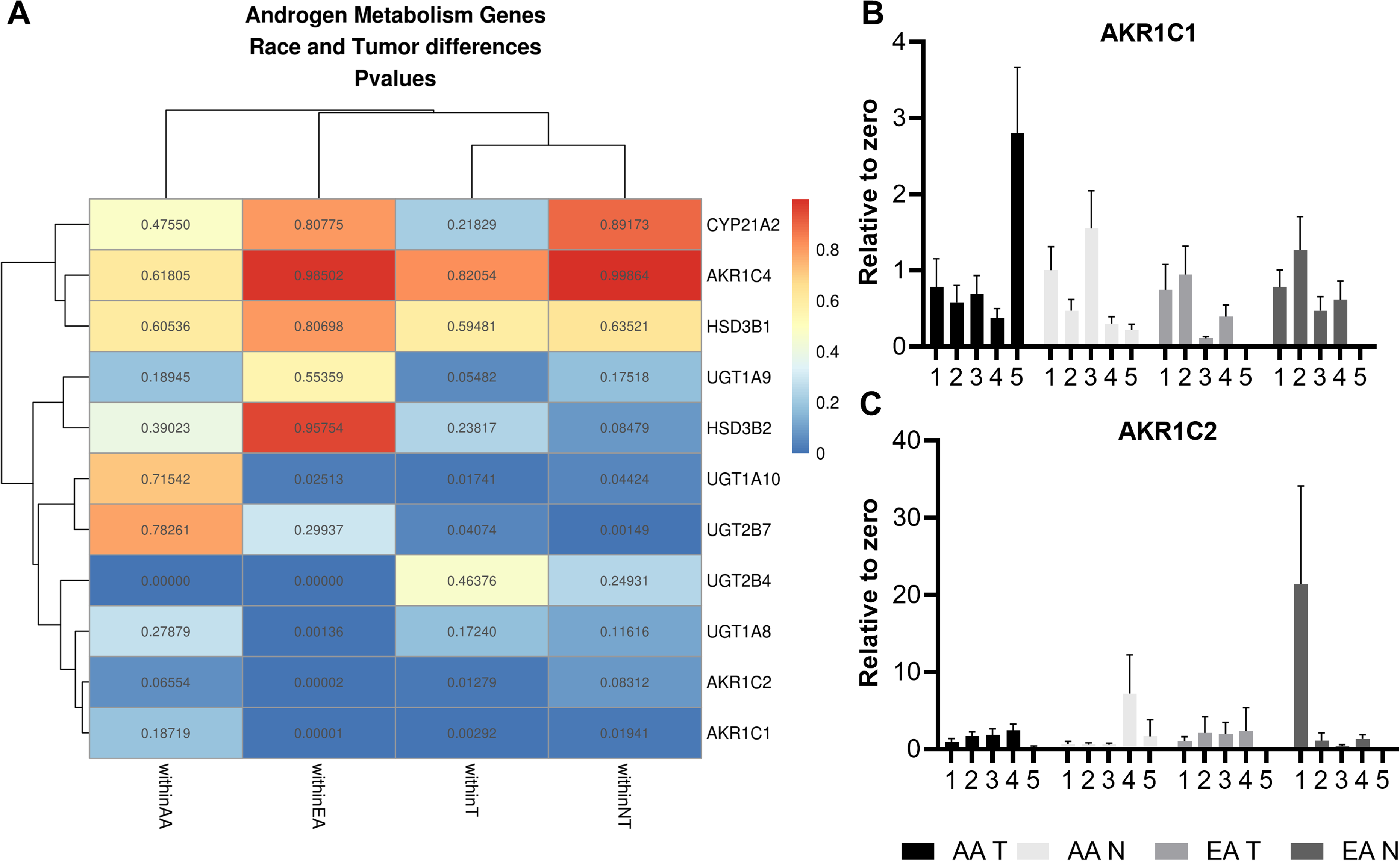
Enzymes involved in DHT synthesis from alternate pathways are significantly dysregulated in PrCa and adjacent non-tumor tissues from AA and EA men. **A.** The heatmap represents the computed p-values obtained upon race and tissue comparisons of enzymes involved in alternate pathways of androgen synthesis. Two-group test p-values within factors were performed for race comparisons. Within race, paired sample t-test is performed, accounting for the paired structure of the data. Each column represents comparisons between tumor and adjacent non-tumor tissues or tissue-specific comparisons between AA and EA men. Dark blue rectangles represent those comparisons that are significantly different in each comparison. **B and C.** qPCR validation of RNA-sequencing results show differential expression of AKR1C1 and AKR1C2 in a subset of PrCa and adjacent non-tumor tissues from AA and EA men.

We applied a metabolism-specific analysis pipeline to Roswell Park Cohort 1^47^. We found that AKR1C, UGT1, and SRD5A gene families were significantly different between PrCa from AA and EA men (**Supplementary Table 5-6**). These genes were included in the gene ontologies of drug metabolism by other enzymes and steroid hormone biosynthesis. We extended the metabolism-specific pipeline to The Cancer Genome Atlas dataset (TCGA) (AA n=43, EA n=270) (**Supplementary Table 5-6**). This discovery pipeline revealed race-specific differences in metabolic pathways, such as tryptophan metabolism, norepinephrine, and purine biosynthesis in the Roswell Park Cohort 1 and TCGA PrCa database (**Supplementary Table 5-6**). Our results suggested that androgen metabolism and other metabolic pathways are already dysregulated at the time of RP in PrCa and adjacent non-tumor tissues in AA compared to EA men.

### Serum DHT levels and expression of androgen metabolizing enzymes differ between AA and EA men with PrCa

The upregulation of RNAs encoding enzymes involved in alternate pathways of androgen synthesis in AA men led us to hypothesize that these may reflect differences in levels of the AR binding androgens, DHT and T. We measured DHT and T in serum since they are synthesized in the different sites, including the Leydig cells of the testis, adrenal glands and tumor/adjacent non-tumor tissues, and freely circulate in the serum^48^. Furthermore, these measurements provide us with a systemic view of androgen levels in AA and EA men with PrCa. We used high-pressure liquid chromatographic assays with tandem mass spectral detection (LC-MS/MS)^49^ to measure levels of T and DHT in the serum of AA (n=38) and EA (n=69) men with PrCa treated at Roswell Park referred to as Roswell Park Cohort 2 (**Supplementary Table 1**, Demographics in **Supplementary Table 7**). The calibrations and performance data are listed in **Supplementary data 8 and 9**. With the exception of one clinical sample, all the serum samples were obtained from patients that received no prior treatments. The median levels of T in the serum were significantly different (p<0.05) with lower levels of T more commonly found in AA compared to EA men with PrCa in Roswell Park Cohort 2 (**Figure 3**, **Table 1**). The median levels of DHT in the serum Roswell Park Cohort 2 was similar between the two racial groups (**Figure 3**, **Table 1**). In the pooled analyses of Roswell Park Cohort 2 that includes both AA and EA men, the median levels of T (p<0.01) and DHT (p=0.053) in the serum were lower in men with PrCa >= Gleason score 7 compared to men with PrCa Gleason score <7 (**Supplementary Table 10**).

**Figure 3.**
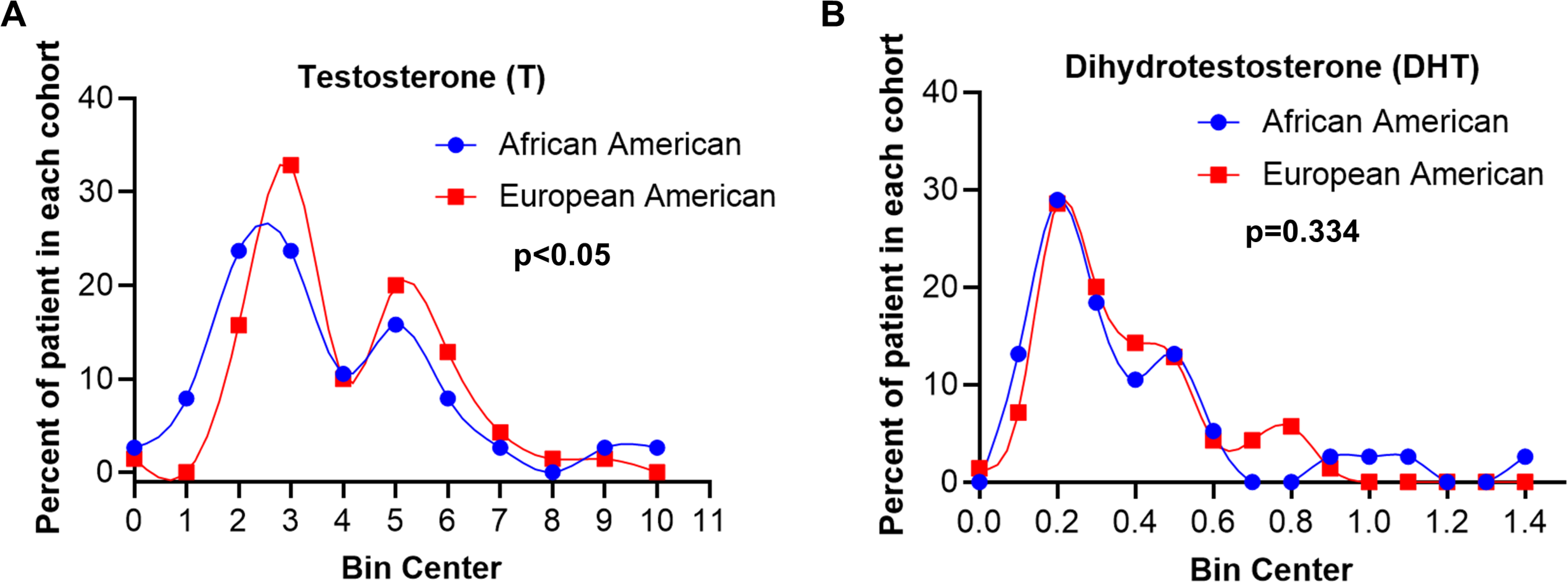
Distribution of T and DHT in the serum of AA and EA men with PrCa. **A.** LC-MS/MS analysis of androgens reveals that the distribution of T levels are significantly different in the serum of AA men compared to EA men with PrCa (p<0.05). **B.** The distribution of DHT in the serum are comparable between AA men and EA men with PrCa. Blue lines represent the frequency of distribution of T and DHT levels in the serum of AA men with PrCa, and pink lines represent the frequency of distribution of T and DHT levels in the serum of EA men with PrCa.

**Table 1.**
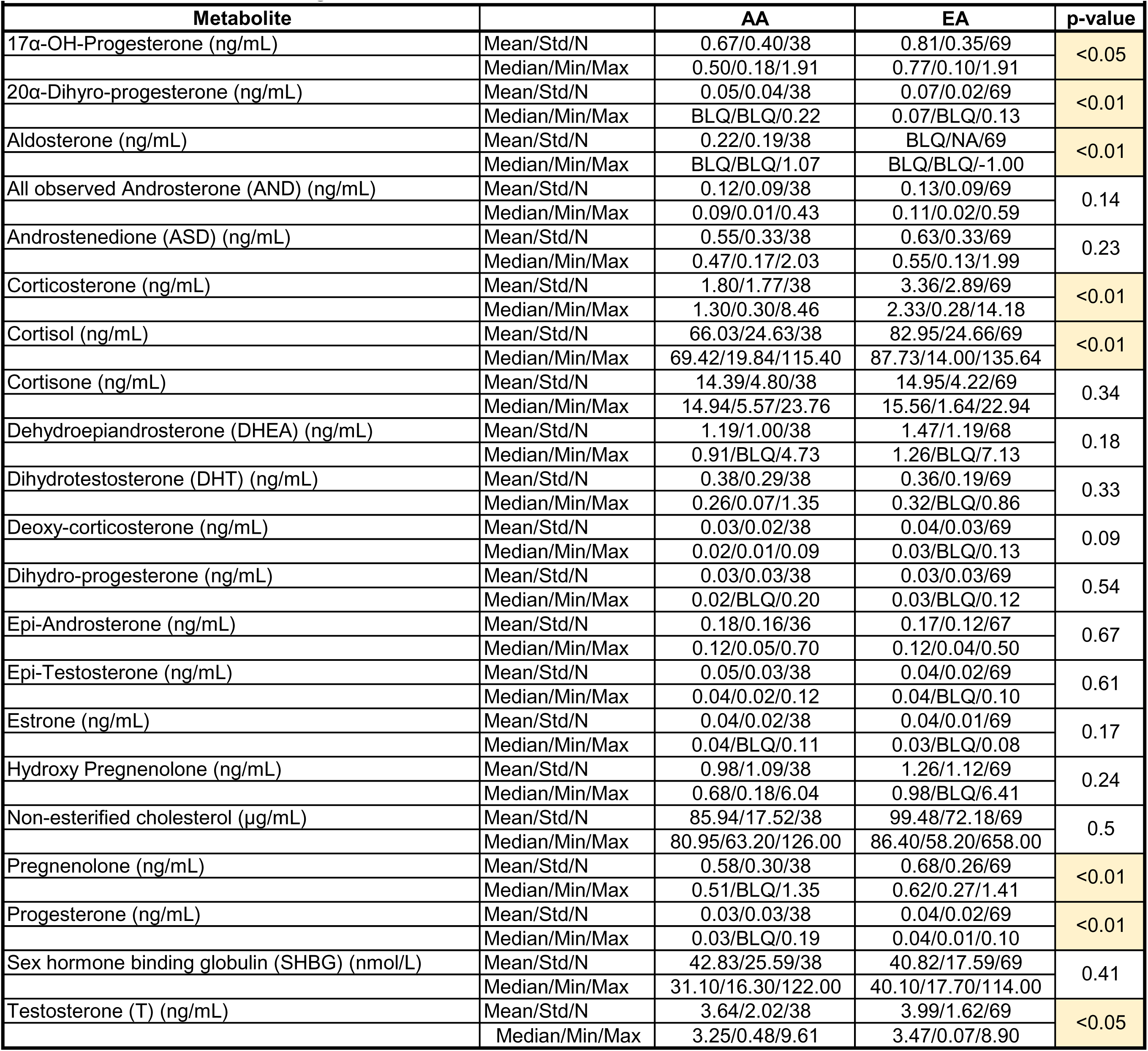
Androgen metabolites measured in the serum of AA and EA men with PrCa

| Metabolite |  | AA | EA | p-value |
| --- | --- | --- | --- | --- |
| 17α-OH-Progesterone (ng/mL) | Mean/Std/N | 0.67/0.40/38 | 0.81/0.35/69 | <0.05 |
|  | Median/Min/Max | 0.50/0.18/1.91 | 0.77/0.10/1.91 |  |
| 20α-Dihydro-progesterone (ng/mL) | Mean/Std/N | 0.05/0.04/38 | 0.07/0.02/69 | <0.01 |
|  | Median/Min/Max | BLQ/BLQ/0.22 | 0.07/BLQ/0.13 |  |
| Aldosterone (ng/mL) | Mean/Std/N | 0.22/0.19/38 | BLQ/NA/69 | <0.01 |
|  | Median/Min/Max | BLQ/BLQ/1.07 | BLQ/BLQ/-1.00 |  |
| All observed Androsterone (AND) (ng/mL) | Mean/Std/N | 0.12/0.09/38 | 0.13/0.09/69 | 0.14 |
|  | Median/Min/Max | 0.09/0.01/0.43 | 0.11/0.02/0.59 |  |
| Androstenedione (ASD) (ng/mL) | Mean/Std/N | 0.55/0.33/38 | 0.63/0.33/69 | 0.23 |
|  | Median/Min/Max | 0.47/0.17/2.03 | 0.55/0.13/1.99 |  |
| Corticosterone (ng/mL) | Mean/Std/N | 1.80/1.77/38 | 3.36/2.89/69 | <0.01 |
|  | Median/Min/Max | 1.30/0.30/8.46 | 2.33/0.28/14.18 |  |
| Cortisol (ng/mL) | Mean/Std/N | 66.03/24.63/38 | 82.95/24.66/69 | <0.01 |
|  | Median/Min/Max | 69.42/19.84/115.40 | 87.73/14.00/135.64 |  |
| Cortisone (ng/mL) | Mean/Std/N | 14.39/4.80/38 | 14.95/4.22/69 | 0.34 |
|  | Median/Min/Max | 14.94/5.57/23.76 | 15.56/1.64/22.94 |  |
| Dehydroepiandrosterone (DHEA) (ng/mL) | Mean/Std/N | 1.19/1.00/38 | 1.47/1.19/68 | 0.18 |
|  | Median/Min/Max | 0.91/BLQ/4.73 | 1.26/BLQ/7.13 |  |
| Dihydrotestosterone (DHT) (ng/mL) | Mean/Std/N | 0.38/0.29/38 | 0.36/0.19/69 | 0.33 |
|  | Median/Min/Max | 0.26/0.07/1.35 | 0.32/BLQ/0.86 |  |
| Deoxy-corticosterone (ng/mL) | Mean/Std/N | 0.03/0.02/38 | 0.04/0.03/69 | 0.09 |
|  | Median/Min/Max | 0.02/0.01/0.09 | 0.03/BLQ/0.13 |  |
| Dihydro-progesterone (ng/mL) | Mean/Std/N | 0.03/0.03/38 | 0.03/0.03/69 | 0.54 |
|  | Median/Min/Max | 0.02/BLQ/0.20 | 0.03/BLQ/0.12 |  |
| Epi-Androsterone (ng/mL) | Mean/Std/N | 0.18/0.16/36 | 0.17/0.12/67 | 0.67 |
|  | Median/Min/Max | 0.12/0.05/0.70 | 0.12/0.04/0.50 |  |
| Epi-Testosterone (ng/mL) | Mean/Std/N | 0.05/0.03/38 | 0.04/0.02/69 | 0.61 |
|  | Median/Min/Max | 0.04/0.02/0.12 | 0.04/BLQ/0.10 |  |
| Estrone (ng/mL) | Mean/Std/N | 0.04/0.02/38 | 0.04/0.01/69 | 0.17 |
|  | Median/Min/Max | 0.04/BLQ/0.11 | 0.03/BLQ/0.08 |  |
| Hydroxy Pregnenolone (ng/mL) | Mean/Std/N | 0.98/1.09/38 | 1.26/1.12/69 | 0.24 |
|  | Median/Min/Max | 0.68/0.18/6.04 | 0.98/BLQ/6.41 |  |
| Non-esterified cholesterol (μg/mL) | Mean/Std/N | 85.94/17.52/38 | 99.48/72.18/69 | 0.5 |
|  | Median/Min/Max | 80.95/63.20/126.00 | 86.40/58.20/658.00 |  |
| Pregnenolone (ng/mL) | Mean/Std/N | 0.58/0.30/38 | 0.68/0.26/69 | <0.01 |
|  | Median/Min/Max | 0.51/BLQ/1.35 | 0.62/0.27/1.41 |  |
| Progesterone (ng/mL) | Mean/Std/N | 0.03/0.03/38 | 0.04/0.02/69 | <0.01 |
|  | Median/Min/Max | 0.03/BLQ/0.19 | 0.04/0.01/0.10 |  |
| Sex hormone binding globulin (SHBG) (nmol/L) | Mean/Std/N | 42.83/25.59/38 | 40.82/17.59/69 | 0.41 |
|  | Median/Min/Max | 31.10/16.30/122.00 | 40.10/17.70/114.00 |  |
| Testosterone (T) (ng/mL) | Mean/Std/N | 3.64/2.02/38 | 3.99/1.62/69 | <0.05 |
|  | Median/Min/Max | 3.25/0.48/9.61 | 3.47/0.07/8.90 |  |

Next, we determined if DHT levels in the serum are indicative of tissue transcript levels of upregulated enzyme encoding genes involved in alternate pathways of DHT biosynthesis. We developed scores using the Gene Set Variation Analysis (GSVA) for gene expression of multiple enzymes involved in alternate pathways, namely *AKR1C3*, *DHRS9*, *HSD17B3*, *HSD3B1*, *HSD3B2*, *RDH5*, *SRD5A1*, *SRD5A2* and *CYP19A1*, for both PrCa and adjacent non-tumor tissues from AA and EA men. We performed these analyses on overlapping samples between Roswell Park Cohorts 1 and 2 (**Supplementary Table 1**). Significantly, we found that DHT levels in the serum were negatively correlated with GSVA score in adjacent non-tumor tissues (r=-0.86, p<0.05) only in AA men (**Figure 4**) but not in EA men with PrCa (**Figure 4**). However, when gene expression in PrCa tissues is normalized to gene expression in adjacent non-tumor tissue, we found a trend towards a positive correlation between genes involved in the alternate pathway for DHT synthesis and DHT levels in the serum (r=0.62) (**Figure 4**). Furthermore, using the Fisher’s r to z conversion we found that there is a significant difference between AA and EA men in correlation patterns between DHT levels in the serum and expression of enzymes involved in the alternate pathway for DHT synthesis in PrCa and adjacent non-tumor tissues (p<0.05, **Figure 4, boxed p-values**). Overall, our results indicate that DHT levels in the serum can potentially be used as surrogate markers for the expression of enzymes involved in alternate pathways for DHT biosynthesis in PrCa and adjacent non-tumor tissue from AA men.

**Figure 4.**
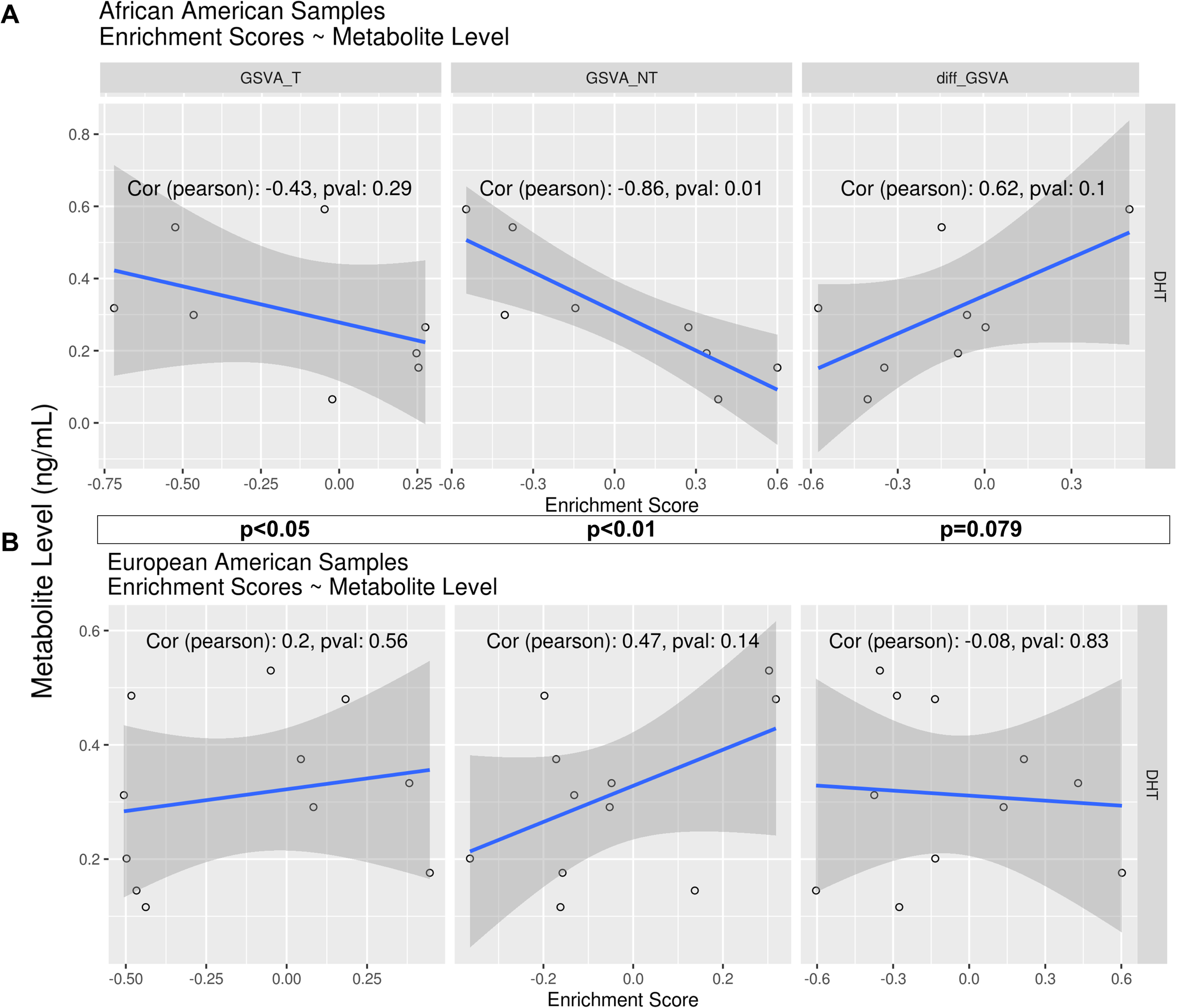
Expression of enzymes involved in DHT biosynthesis from alternate pathways are correlated to DHT levels in the serum of AA men with PrCa. GSVA scores were calculated for the following enzymes: *AKR1C3*, *DHRS9*, *HSD17B3*, *HSD3B1*, *HSD3B2*, *RDH5*, *SRD5A1*, *SRD5A2,* and *CYP19A1*. Pearson correlation analysis were calculated for GSVA scores and DHT levels in the serum. The first column displays correlation with expression in PrCa tissues (T). The second column displays correlation with expression in adjacent non-tumor tissues (NT). The third column displays correlation with normalized expression in tumor tissues (T/NT). The X-axis represents the GSVA score, and the Y-axis represents the bin centers from frequency distribution analysis of DHT levels in the serum (ng/mL). **A.** Represents clinical samples from AA men with PrCa **B.** Represents clinical samples from EA men with PrCa.

### Alternate DHT biosynthesis pathways are used by AA men at PrCa diagnosis

Since the median levels of T but not DHT in the serum of AA men with PrCa was significantly lower compared to EA men with PrCa (**Figure 3**, **Table 1**), we investigated if this was potentially due to differences in pathways utilized for DHT biosynthesis. For this purpose, we analyzed levels of androgen intermediate metabolites from the different pathways in the serum of AA and EA men with PrCa. We first determined the serum levels of cholesterol, an important precursor for Leydig/adrenal as well as *de novo* tumor T and DHT synthesis^4^. The median levels of non-esterified cholesterol in the serum was similar between AA (n=38) and EA (n=69) men with PrCa in Roswell Park Cohort 2 (**Table 1**). This was consistent with our RNA-sequencing analyses of Roswell Park Cohort 1 that did not detect differences in the expression of enzymes that are essential for cholesterol synthesis, including Farnesyl-Diphosphate Farnesyltransferase 1 (FDFT1) and Squalene monooxygenase (SQLE), in PrCa and adjacent non-tumor tissues from AA and EA men (**Supplementary Figure 2**). This observation combined with the correlations between DHT and the expression of enzyme genes in the alternative pathways also provided confidence in measuring levels in the serum as surrogates for tissue production of androgen metabolites.

Significant differences in T but not non-esterified cholesterol and DHT levels in the serum of AA and EA men can possibly reflect the choice of pathways downstream from cholesterol utilized for DHT synthesis. Alternatively, it is possible that DHT might not be further converted to another downstream metabolite. We tested the first scenario by comprehensively measuring the levels of intermediate metabolites produced during DHT and T biosynthesis^1, 29, 30^ in the serum of AA (n=38) and EA (n=69) men with PrCa in Roswell Park Cohort 2 (**Supplementary Table 1**). The grey highlighted box in **Supplementary Figure 3** depicts the primary androgen synthesis pathway^1, 29, 30^. Although we can’t be certain of where these intermediate metabolites are produced, AA men with PrCa had an increased frequency of low levels of pregnenolone in the serum compared to EA men with PrCa (65.8 vs. 42%, p<0.05) (**Supplementary Table 11**). This was reflected in the median levels of pregnenolone which was significantly lower in the serum of AA compared to EA men with PrCa (p<0.01) (**Table 1**). Additionally, AA men with PrCa were more likely to have low levels of hydroxy pregnenolone (63.2 vs 43.5%, p=0.069) and DHEA (63 vs 44%, p=0.071), compared to EA men with PrCa (**Table 1, Supplementary Table 11**). The median levels of ASD, a final and penultimate metabolite before conversion to T and DHT, respectively^1, 29, 30^, was lower, but not significant, in the serum of AA compared to EA men with PrCa (**Table 1, Supplementary Table 11**). Our data demonstrate that AA men with PrCa exhibit lower serum levels of intermediate metabolites generated from the primary pathway for DHT and T synthesis.

We next measured the levels of intermediate metabolites produced through the alternate pathways of DHT and T biosynthesis in the serum of AA and EA men with PrCa. Pregnenolone can be converted to T and DHT through an intermediate progesterone conversion step^1, 29, 30^. Significantly, the median levels of progesterone (p<0.01) and its associated metabolites, including 17α-OH-progesterone (p<0.05), cortisol (p<0.01), 20α-dihydro-progesterone (p<0.01) and corticosterone (p<0.01) were lower in AA men compared to EA men with PrCa in Roswell Park Cohort 2 (**Figure 5**, **Table 1**). Progesterone can be converted to ASD and to AND that leads to DHT synthesis through 5α-dione (**Supplementary Figure 3**). The median levels of AND in the serum were similar between AA men and EA men with PrCa (**Table 1**). Significantly, AND levels were positively correlated to DHT levels only in the serum of AA men with PrCa (**Supplementary Figure 4**). These data suggest that DHT levels in the serum can be a surrogate indicator of AND levels in the serum of AA men with PrCa. AND can be converted to epiandrosterone that is further sulfated to act as a reserve for DHT production^50^. The median levels of epiandrosterone in the serum were similar between the two racial groups (**Table 1**). Overall, we find lower levels of progesterone related metabolites in AA men with PrCa and similar levels of AND in AA and EA men with PrCa. These data provide evidence that AA men with PrCa are more likely to use an alternate progesterone pathway for DHT biosynthesis.

**Figure 5.**
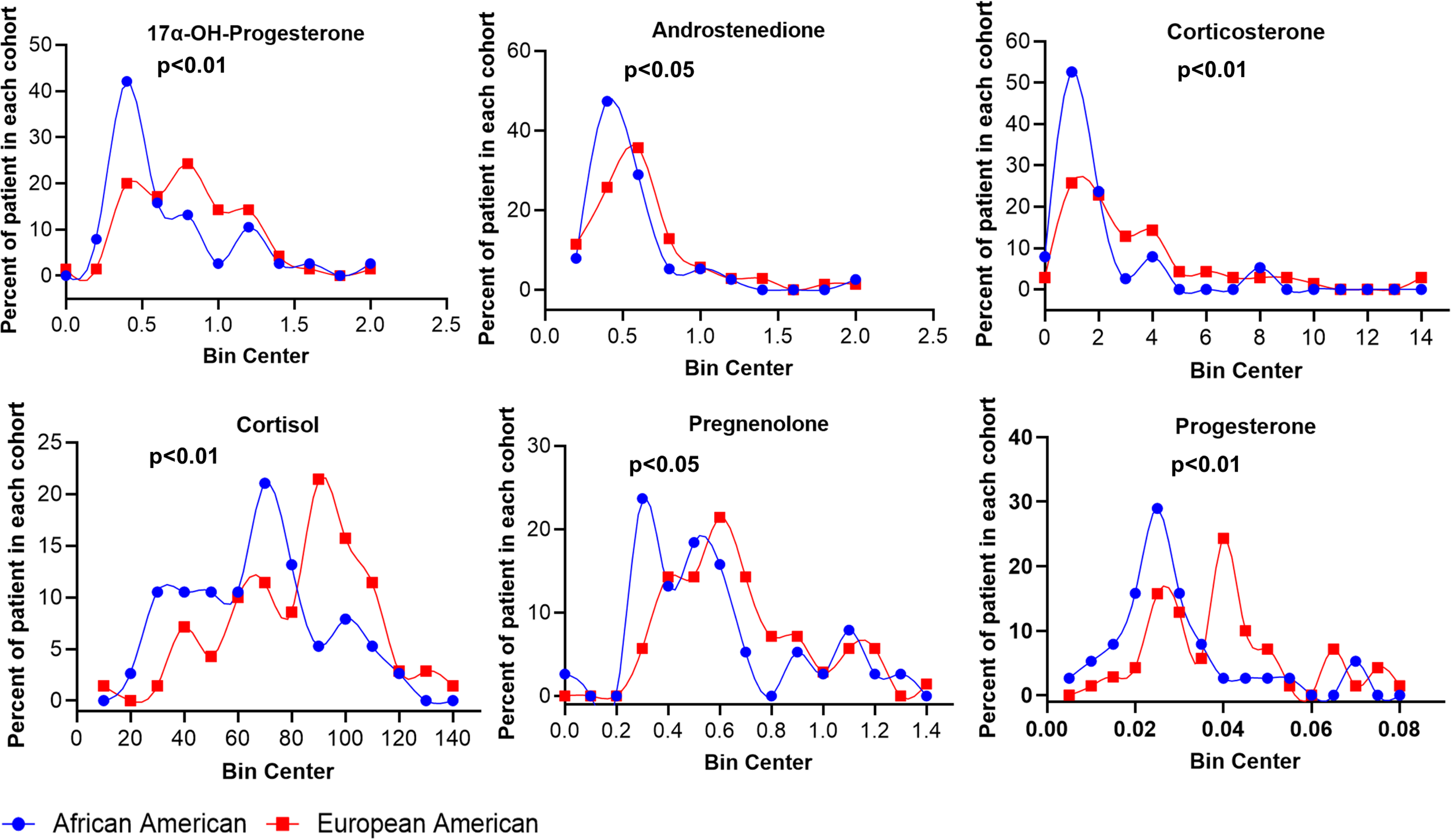
Serum distribution of intermediate metabolites produced during alternate DHT biosynthesis pathways in AA and EA men with PrCa. LC-MS/MS analysis reveals that the frequency of high and low intermediate androgen metabolites levels in the serum are significantly different between AA men and EA men with PrCa.

### Serum aldosterone levels are associated with West African ancestry markers

Aldosterone is one of the terminal metabolites in the progesterone-corticosterone biosynthesis pathway. We found that aldosterone (p<0.01) levels in the serum were above the limit of detection in a subset of AA men with PrCa in Roswell Park Cohort 1 (**Table 1**). Significantly, we did not detect aldosterone in the any of the serum samples in the Roswell Park Cohort 2 of EA men with PrCa (**Table 1**). Surprisingly, we found that aldosterone levels in the serum positively correlated with AIMs associated with West African ancestry (**Figure 6A**). We initially performed ancestry informative markers (AIMs) analysis to determine concordance between self-reported race and AIMs for a subset of men referred to as Roswell Park Cohort 3 (**Supplementary Table 1**). There was a direct correlation between self-reported race and AIMs in most of the clinical samples in Roswell Park Cohort 3 (**Supplementary Table 1 and 12**). With the observation of positive correlation with AIMs and aldosterone levels in the serum, we next determined if there are any differences in CYP11B enzymes that regulate corticosterone to aldosterone conversion^51^. Examination of exome sequencing analyses in a separate set of AA and EA men with PrCa (Roswell Park Cohort 4, **Supplementary Table 1**) revealed that AA men with PrCa had a significantly higher frequency of multiple single nucleotide polymorphisms (SNPs) in the genes encoding CYP11B1 and CYP11B2 (**Figure 6B-C**, **Supplementary Table 13**). Furthermore, two SNPs, rs5313 and rs4544, located in the exon region of CYP11B2 have the potential to affect CYP11B2 function (**Figure 6B-C**, **Supplementary Table 13**). Therefore, it is possible that AA men with SNPs in the CYP11 family of enzymes may more readily convert corticosterone to aldosterone. Limited numbers of overlapping samples between Roswell Park Cohorts 2 and 4 prevented such correlation analyses. Overall, our data indicates that AA men with PrCa can have increased frequency of certain SNPs that can alter aldosterone synthesis.

**Figure 6.**
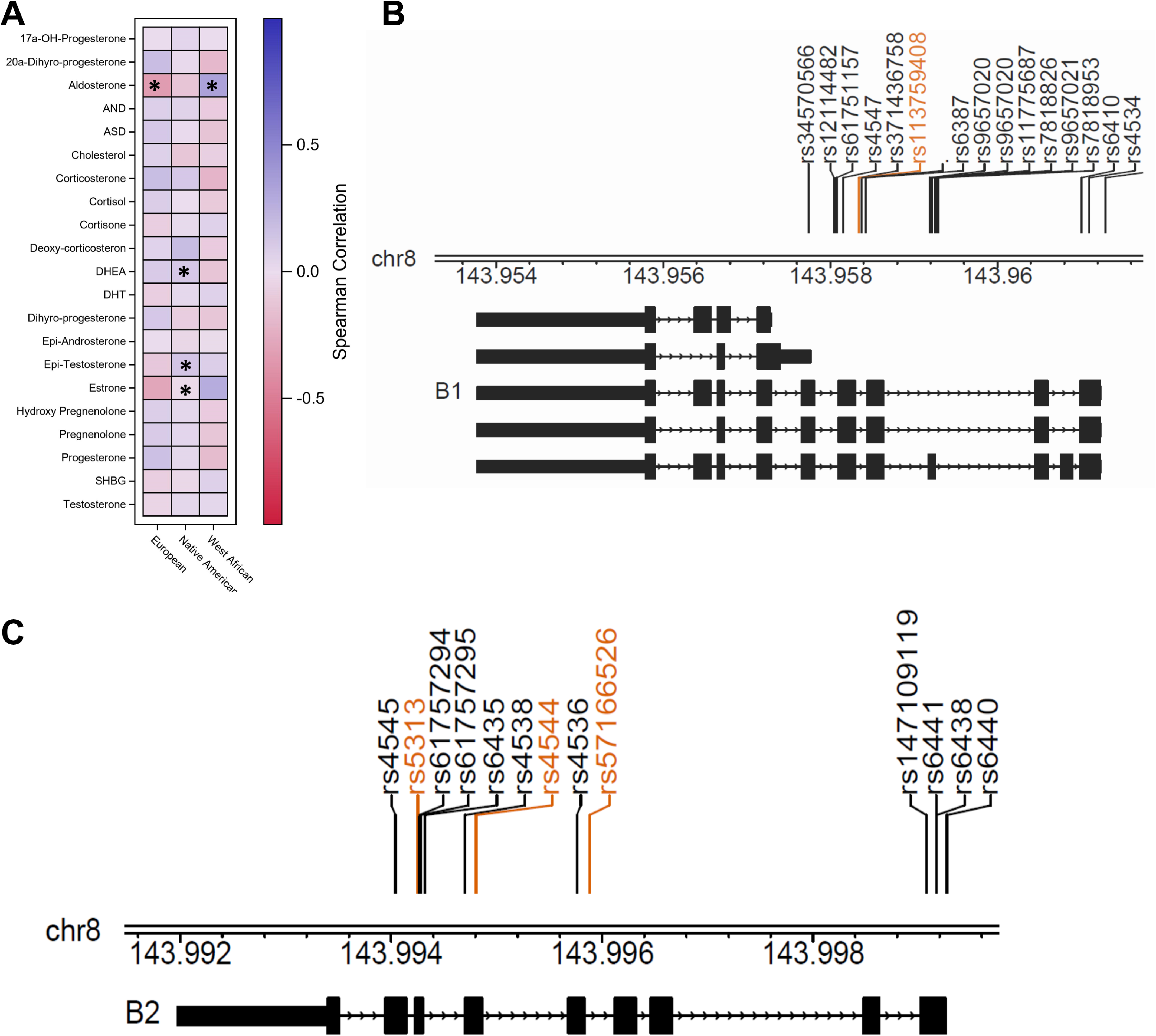
Subset of intermediate metabolites in androgen synthesis measured in the serum are associated with AIMs. **A.** Spearman correlation analysis reveals that a subset of intermediate metabolites is correlated to AIMs. Blue represents positive correlation, and red represents negative correlation. *indicates significant correlation with p-value less than 0.05. **B and C.** The figures represent the SNPs identified in CYP11B1 and CYP11B2 in AA and EA men with PrCa. The SNPs labeled in orange represent those SNPs found at significantly higher frequencies in AA men with PrCa.

### Serum androgen and metabolites are indicators of progression-free survival and RP failure in a race-specific manner

It is unknown whether intermediate metabolites produced during DHT and T synthesis are indicators of clinical outcomes in PrCa. Therefore, we investigated whether levels of DHT/T and related intermediate metabolites in the serum (high vs. low) can be indicators of early disease-progression and whether this is a race-specific indicator in Roswell Park Cohort 2. Overall survival, progression-free survival (PFS) and RP failure (RPF) were similar between AA and EA men with PrCa in Roswell Park Cohort 2 (**Supplementary Figure 5**). RPF is defined as the time from surgery until persistent disease, disease progression, subsequent treatment, death from PrCa, or last follow-up. The metabolite levels were dichotomized into low (< 50^th^ percentile) and high (> 50^th^ percentile) based on the median value of the pooled samples or into ‘below lower limit of quantitation’ (BLQ) and detectable (i.e. >BLQ). Since the levels of each metabolite in the serum varied, each metabolite had a different cut-off for dichotomization for survival analyses. Both AA and EA men with PrCa had an equal distribution of high and low levels of non-esterified cholesterol in the serum (**Table 1, Supplementary Table 11**). We found that low levels of non-esterified cholesterol were associated with an increased incidence of 3- (p<0.01) and 5-years RPF (p<0.05) in pooled analyses of both AA and EA men with PrCa (**Figure 7, Supplementary Table 14**). Thus, non-esterified cholesterol levels in the serum are a potential indicators of early disease progression in a race-independent manner. AA men with PrCa more frequently had lower levels of ASD in the serum compared to EA men (65.8 vs. 42%, p<0.05) (**Supplementary Table 11**). In our pooled analyses, men with PrCa and low levels of ASD in the serum had increased incidence of 3-year RPF compared to men with PrCa and high levels of ASD (**Figure 7, Supplementary Table 14**). In the progesterone-cortisone pathway, AA men with PrCa had increased frequency of low levels of 17α-OH-progesterone (68.4 vs. 40.6%, p<0.01) and progesterone (71.1 vs. 39.1%, p<0.01) in the serum compared to EA men with PrCa (**Supplementary Table 11**). Men with low levels of 17α-OH-progesterone in the serum had significantly reduced 3-year, 5-year, and long-term RPF (p<0.05) compared to men with high levels of 17α-OH-progesterone (**Figure 7, Supplementary Table 14**). Similarly, men with low levels of progesterone had lower 5-year (p<0.05) and long term RPF (p=0.06) compared to men with high levels of progesterone (**Figure 7, Supplementary Table 14**). Additionally, low levels of other progesterone-derived metabolites, deoxycorticosterone and corticosterone in the serum, were both indicators of low 5-year RPF in a race-independent manner (**Figure 7, Supplementary Table 14**). Overall, our results indicate that low levels of a subset of intermediate metabolites generated during DHT and T synthesis are found more commonly in AA men with PrCa and could be prognostic indicators of early disease progression.

**Figure 7.**
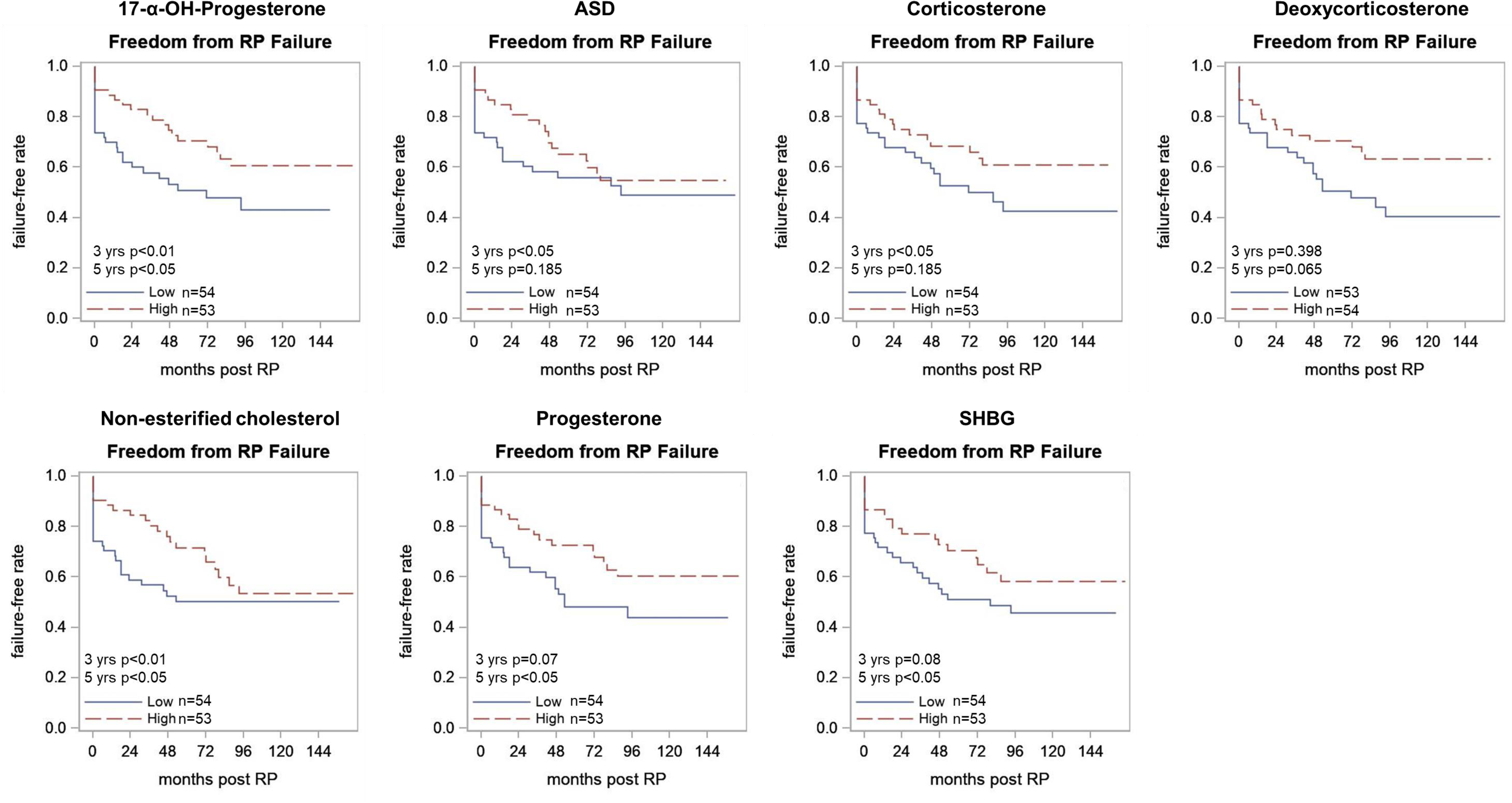
A subset of intermediate metabolites produced during DHT biosynthesis reflects shorter time to disease progression. Lower levels of intermediate androgen metabolites are associated with shorter time to disease progression in Roswell Park Cohort 2.

### Androgen receptor expression and activity is associated with DHT levels and PrCa tumor stage in AA men

Since we observed lower levels of T but not DHT in the serum in AA men with PrCa, we determined whether these levels are correlated with AR expression and transcriptional activity in both PrCa and adjacent non-tumor tissue of AA and EA men. First, we determined the percentage of positive AR nuclei in PrCa and adjacent non-tumor tissues, and their relationship to clinicopathological parameters in AA (n=92) and EA (n=95) men in tissue microarrays (TMAs) Roswell Park Cohort 5 (**Supplementary Table 1**, Demographics in **Supplementary Table 15**). In agreement with previous results^52^, the percentage of positive AR nuclei was significantly higher in adjacent non-tumor tissues (**Supplementary Figure 6**), but not different between tumor tissues, from AA compared to EA men (mean: 78.20 vs. 73.28, p<0.01) (**Table 2, Supplementary Figure 6**). In the pooled analyses of Roswell Park Cohort 5, no significant differences in the percentage of positive AR nuclei in PrCa tissue or non-tumor tissue adjacent to PrCa between PrCa with Gleason scores <=3+4 and >=4+3 were detected (**Table 2**). However, when race specific differences were determined, we observed that the percentage of positive AR nuclei were lower in AA than EA men (mean: 75.19 vs. 78.82, p=0.07) in PrCa with Gleason score >= 4+3 (**Table 2**). By contrast, the percentage of positive AR nuclei was higher in adjacent non-tumor tissues in AA compared to EA men with Gleason score >= 4+3 (mean: 80.15 vs 71.31) (**Table 2**). Surprisingly, AA men who did not receive hormone therapies had a significantly higher percentage of positive AR nuclei compared to AA men who received hormone therapies (80.45 vs 68.78, p<0.001) (**Table 2**). Combined, our data suggest that AR protein expression in both adjacent non-tumor and PrCa tissues are indicators of higher Gleason score only in AA men with PrCa.

**Table 2.**
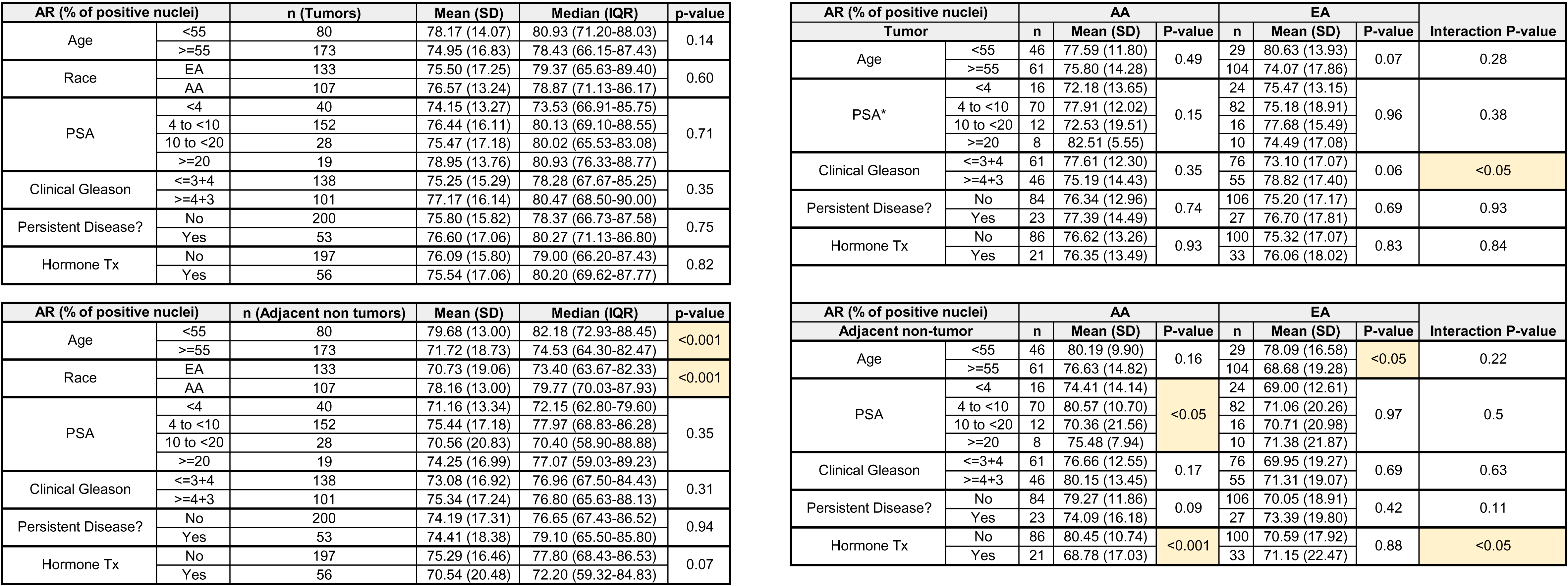
AR protein expression correlates to clinicopathological parameters

| AR (% of positive nuclei) |  | n (Tumors) | Mean (SD) | Median (IQR) | p-value |
| --- | --- | --- | --- | --- | --- |
| Age | <55 | 80 | 78.17 (14.07) | 80.93 (71.20-88.03) | 0.14 |
|  | >=55 | 173 | 74.95 (16.83) | 78.43 (66.15-87.43) |  |
| Race | EA | 133 | 75.50 (17.25) | 79.37 (65.63-89.40) | 0.60 |
|  | AA | 107 | 76.57 (13.24) | 78.87 (71.13-86.17) |  |
| PSA | <4 | 40 | 74.15 (13.27) | 73.53 (66.91-85.75) | 0.71 |
|  | 4 to <10 | 152 | 76.44 (16.11) | 80.13 (69.10-88.55) |  |
|  | 10 to <20 | 28 | 75.47 (17.18) | 80.02 (65.53-83.08) |  |
|  | >=20 | 19 | 78.95 (13.76) | 80.93 (76.33-88.77) |  |
| Clinical Gleason | <=3+4 | 138 | 75.25 (15.29) | 78.28 (67.67-85.25) | 0.35 |
|  | >=4+3 | 101 | 77.17 (16.14) | 80.47 (68.50-90.00) |  |
| Persistent Disease? | No | 200 | 75.80 (15.82) | 78.37 (66.73-87.58) | 0.75 |
|  | Yes | 53 | 76.60 (17.06) | 80.27 (71.13-86.80) |  |
| Hormone Tx | No | 197 | 76.09 (15.80) | 79.00 (66.20-87.43) | 0.82 |
|  | Yes | 56 | 75.54 (17.06) | 80.20 (69.62-87.77) |  |

| AR (% of positive nuclei) |  | n (Adjacent non tumors) | Mean (SD) | Median (IQR) | p-value |
| --- | --- | --- | --- | --- | --- |
| Age | <55 | 80 | 79.68 (13.00) | 82.18 (72.93-88.45) | <0.001 |
|  | >=55 | 173 | 71.72 (18.73) | 74.53 (64.30-82.47) |  |
| Race | EA | 133 | 70.73 (19.06) | 73.40 (63.67-82.33) | <0.001 |
|  | AA | 107 | 78.16 (13.00) | 79.77 (70.03-87.93) |  |
| PSA | <4 | 40 | 71.16 (13.34) | 72.15 (62.80-79.60) | 0.35 |
|  | 4 to <10 | 152 | 75.44 (17.18) | 77.97 (68.83-86.28) |  |
|  | 10 to <20 | 28 | 70.56 (20.83) | 70.40 (58.90-88.88) |  |
|  | >=20 | 19 | 74.25 (16.99) | 77.07 (59.03-89.23) |  |
| Clinical Gleason | <=3+4 | 138 | 73.08 (16.92) | 76.96 (67.50-84.43) | 0.31 |
|  | >=4+3 | 101 | 75.34 (17.24) | 76.80 (65.63-88.13) |  |
| Persistent Disease? | No | 200 | 74.19 (17.31) | 76.65 (67.43-86.52) | 0.94 |
|  | Yes | 53 | 74.41 (18.38) | 79.10 (65.50-85.80) |  |
| Hormone Tx | No | 197 | 75.29 (16.46) | 77.80 (68.43-86.53) | 0.07 |
|  | Yes | 56 | 70.54 (20.48) | 72.20 (59.32-84.83) |  |

| AR (% of positive nuclei) |  | AA |  |  | EA |  |  |  |
| --- | --- | --- | --- | --- | --- | --- | --- | --- |
| Tumor |  | n | Mean (SD) | P-value | n | Mean (SD) | P-value | Interaction P-value |
| Age | <55 | 46 | 77.59 (11.80) | 0.49 | 29 | 80.63 (13.93) | 0.07 | 0.28 |
|  | >=55 | 61 | 75.80 (14.28) |  | 104 | 74.07 (17.86) |  |  |
| PSA* | <4 | 16 | 72.18 (13.65) | 0.15 | 24 | 75.47 (13.15) | 0.96 | 0.38 |
|  | 4 to <10 | 70 | 77.91 (12.02) |  | 82 | 75.18 (18.91) |  |  |
|  | 10 to <20 | 12 | 72.53 (19.51) |  | 16 | 77.68 (15.49) |  |  |
|  | >=20 | 8 | 82.51 (5.55) |  | 10 | 74.49 (17.08) |  |  |
| Clinical Gleason | <=3+4 | 61 | 77.61 (12.30) | 0.35 | 76 | 73.10 (17.07) | 0.06 | <0.05 |
|  | >=4+3 | 46 | 75.19 (14.43) |  | 55 | 78.82 (17.40) |  |  |
| Persistent Disease? | No | 84 | 76.34 (12.96) | 0.74 | 106 | 75.20 (17.17) | 0.69 | 0.93 |
|  | Yes | 23 | 77.39 (14.49) |  | 27 | 76.70 (17.81) |  |  |
| Hormone Tx | No | 86 | 76.62 (13.26) | 0.93 | 100 | 75.32 (17.07) | 0.83 | 0.84 |
|  | Yes | 21 | 76.35 (13.49) |  | 33 | 76.06 (18.02) |  |  |
| Adjacent non-tumor |  | n | Mean (SD) | P-value | n | Mean (SD) | P-value | Interaction P-value |
| Age | <55 | 46 | 80.19 (9.90) | 0.16 | 29 | 78.09 (16.58) | <0.05 | 0.22 |
|  | >=55 | 61 | 76.63 (14.82) |  | 104 | 68.68 (19.28) |  |  |
| PSA | <4 | 16 | 74.41 (14.14) | <0.05 | 24 | 69.00 (12.61) | 0.97 | 0.5 |
|  | 4 to <10 | 70 | 80.57 (10.70) |  | 82 | 71.06 (20.26) |  |  |
|  | 10 to <20 | 12 | 70.36 (21.56) |  | 16 | 70.71 (20.98) |  |  |
|  | >=20 | 8 | 75.48 (7.94) |  | 10 | 71.38 (21.87) |  |  |
| Clinical Gleason | <=3+4 | 61 | 76.66 (12.55) | 0.17 | 76 | 69.95 (19.27) | 0.69 | 0.63 |
|  | >=4+3 | 46 | 80.15 (13.45) |  | 55 | 71.31 (19.07) |  |  |
| Persistent Disease? | No | 84 | 79.27 (11.86) | 0.09 | 106 | 70.05 (18.91) | 0.42 | 0.11 |
|  | Yes | 23 | 74.09 (16.18) |  | 27 | 73.39 (19.80) |  |  |
| Hormone Tx | No | 86 | 80.45 (10.74) | <0.001 | 100 | 70.59 (17.92) | 0.88 | <0.05 |
|  | Yes | 21 | 68.78 (17.03) |  | 33 | 71.15 (22.47) |  |  |

We next investigated whether the percentage of positive AR nuclei in PrCa was correlated with T and DHT levels in the serum by utilizing a subset of clinical samples that overlapped in Roswell Park Cohorts 2 and 5 (**Supplementary Table 1**). DHT levels in the serum of all men trended towards a negative correlation with percentage of positive AR nuclei in adjacent non-tumor tissues of AA but not EA men (r=-0.21 vs r=0.09) (**Supplementary Figure 7**). However, T levels in the serum trended towards negative association with the percentage of positive AR nuclei in adjacent non-tumor tissues in both AA and EA men with PrCa (r=-0.14 vs r=-0.03) (**Supplementary Figure 7**). In PrCa tissues, the percentage of positive AR nuclei trended towards a negative correlation with DHT and T levels in the serum of both AA and EA men PrCa (**Supplementary Figure 7**). The lower r values in the correlation analyses could be a result of lower availability of overlapping clinical samples for both serum metabolite and AR protein expression analyses, and thus, should be interpreted with caution. Nevertheless, these results underscore the importance of further characterization of androgens and intermediate metabolites in the serum along with AR gene and protein level data in a larger cohort of clinical samples.

Finally, we determined if T and DHT levels in the serum correlate with AR activity in adjacent non-tumor and PrCa tissues in a race-specific manner. We first observed the levels of SHBG, a protein that binds T and DHT to reduce the levels of circulating bioavailable androgens, in the serum of AA and EA men with PrCa^53^. Previous studies have shown that SHBG levels in the serum are variable in AA and EA populations regardless of whether these men are healthy or have PrCa^17, 32, 54–61^. In our study, the median levels of SHBG in the serum were similar between AA and EA men with PrCa (**Table 1**). Moreover, SHBG levels were positively correlated to T and DHT in all clinical samples (**Supplementary Figure 4**). These data suggest that the bioavailable levels of DHT can potentially be similar between AA and EA men. Next, we investigated a subset of patient specimens that were analyzed for both RNA sequencing (Roswell Park Cohort 3) and serum metabolite measurements (Roswell Park Cohort 1) (AA n=8, EA n=11) (**Supplementary Table 1**). For AR activity, we measured RNA levels of the established AR transcriptional targets *KLK2*, *KLK3*, *NKX3.1,* and *TMPRSS2* that have consistently validated as AR transcriptional targets in multiple studies^62–65^. We utilized the GSVA approach to develop a score based on RNA expression of these four target genes. There was insufficient power to compare AR activity between clinical samples with low and high levels of T and DHT, as well as AR protein expression, due to the limited number of overlapping samples between RNA sequencing and serum metabolite analyses. However, we found that DHT levels in the serum correlated with AR activity in AA men, but not EA men, with PrCa (**Figure 8**). DHT levels in the serum of AA men with PrCa trended toward a positive correlation with AR activity in PrCa tissues (**Figure 8**). There was a significant correlation (r=0.75, p<0.05) when RNA expression in PrCa tissues was normalized to RNA expression in adjacent non-tumor tissues (**Figure 8**). Overall, our results indicate that DHT levels in the serum reflected AR transcriptional activity only in PrCa from AA men.

**Figure 8.**
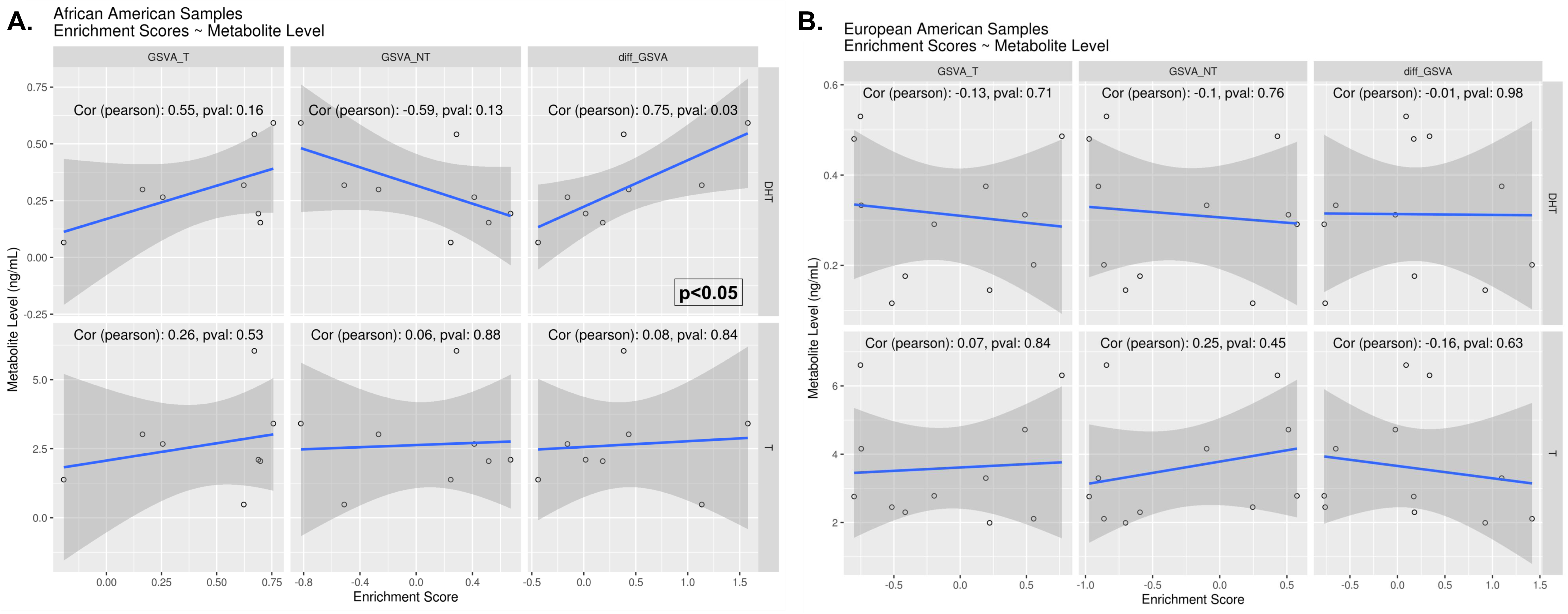
AR transcriptional activity is positively correlated to DHT levels in the serum of AA men with PrCa. GSVA scores were calculated for AR transcriptional targets: *KLK2*, *KLK3*, *TMPRSS2 and NKX3.1,* followed by correlation analysis with androgen levels in the serum. The first column displays correlation with expression in PrCa tissues (T). The second column displays correlation with expression in adjacent non-tumor tissues (NT). The third column displays correlation with normalized expression in tumor tissues (T/NT). The X-axis represents the GSVA score, and the Y-axis represents the level of androgens in the serum (ng/mL). The top panel is the correlation between DHT and AR activity, and the bottom panels represent the correlation between T and AR activity. **A.** Represents clinical samples from AA men with PrCa **B.** Represents clinical samples from EA men with PrCa.

## Discussion

Our studies show that overexpression of enzymes from alternate pathways in androgen synthesis in therapy naïve RP tissues correlates with DHT levels in the serum only in AA men. These data suggest that changes in the androgen synthesis pathway is an early occurrence in PrCa in AA men. It is vital that androgen and intermediate metabolite measurements in the serum are conducted using reliable and sensitive technologies. We used the LC-MS/MS technique^49^, which is sensitive and specific, to identify race-specific differences in the levels of androgens and several intermediate androgen metabolites. The LC-MS/MS approach based on mass and time of elution accurately predicts the presence of a particular androgen related metabolite in the given sample. The use of LC-MS/MS could also be a reason for discrepant results from previously published reports that use relatively less sensitive chemiluminescent assays and radioimmunoassays^18, 45, 66, 67^. Our measurements were performed in serum, although PrCa itself can produce androgens^25, 31, 32, 49, 68^. There are reports of race-specific differences in levels of intermediate androgen metabolites, but neither T nor DHT levels in PrCa tissues in the castration recurrent setting^25, 31, 32, 49, 68^. Future studies that measure the levels of androgens in therapy naïve PrCa tissues and serum from the same clinical cohort will be important to understand the relationship between androgens in circulation and in tissues.

Our pooled data show that lower levels of T and DHT in the serum are associated with high Gleason score PrCa. Roswell Park Cohort 2 has a higher percentage of AA men with PrCa with Gleason score >7 compared to EA men. Therefore, there are two possible explanations for these data. The first is that AA men with PrCa with Gleason score >7 are more likely to have lower serum T levels than EA men with PrCa with Gleason score >7. Another explanation would be that Gleason score is a potential confounding factor. A multivariate analysis is required to address this, but the analysis of a greater number of clinical samples needs to be done to confirm this hypothesis. Our studies also show that mean T levels but not DHT levels in the serum are significantly lower in AA men with PrCa, whereas mean SHBG levels are similar between AA and EA men with PrCa. Based on our findings and what is known about SHBG^48, 69, 70^ and its impact on circulating hormones, bioavailable T in tissues may be even lower in AA compared to EA men with PrCa. A rapid fall in levels of T in the serum which would lower bioavailable T are more commonly observed in healthy AA men, and this rapid fall is associated with PrCa risk^17, 18, 54, 55, 58, 59, 71–73^. It is important to note that this study focused on serum that was obtained at the time of PrCa diagnosis and we did not investigate serum levels of androgens and intermediate metabolites either before disease diagnosis or throughout the disease continuum. An ideal future study would follow the same cohort of patients and assess if there are race-specific associations between levels of DHT and T in serum at multiple points with PrCa progression. It is counterintuitive that rapid falls in T levels during castration and potentially androgen deprivation therapies could contribute to PrCa progression. Whether targeting androgen metabolism earlier in the PrCa continuum would also cause a dramatic fall in DHT/T levels, and therefore lead to rapid disease progression, remains to be investigated.

Interestingly, our pooled analyses revealed that lower levels of several intermediate metabolites were associated with lower time to RPF, thus suggesting these are indicators of early disease progression. Additionally, AA men with PrCa more often present with lower levels of these metabolites in serum. Overall survival, PFS and RPF were similar between AA and EA men in the Roswell Park clinical cohorts. Therefore, we provide a rationale for measuring the intermediate metabolites, in addition to DHT and T, in the serum and possibly PrCa tissues and determining their utility as race-specific biomarkers in a larger cohort of PrCa patients. We also discovered higher frequency of novel germline SNPs in the genes encoding CYP11B family of enzymes in AA men with PrCa. The SNPs we report are either missense variants or variants that can possibly affect the coding sequence of CYP11B2, and ultimately affect its function. The CYP11B family of enzymes mediate corticosterone to aldosterone conversion^51^. We find that only a subset of AA men with PrCa have aldosterone levels that are above the limit of detection in serum. Further studies in larger cohorts of different racial groups are needed to establish the causal relationship between the SNPs observed in CYP11B2-encoding genes and corticosterone-aldosterone conversion. These studies can also determine whether SNPs in CYP11B enzymes can be used as potential race-specific biomarkers in a larger cohort of PrCa patients.

Circulating T and DHT from the serum that either actively or passively diffuse through the tissue can maintain AR nuclear translocation and thus transcriptional activity in both PrCa and adjacent non-tumor tissues^48, 69, 70, 74, 75^. In addition, DHT has a greater affinity and slower rate of dissociation from AR^76–79^. Our data that shows similar levels of DHT in the serum suggest that DHT-mediated AR transcriptional activity may be similar between AA and EA men. Although DHT and T are thought to be the primary ligands for AR, intermediate metabolites produced during androgen synthesis also have some affinity to AR^1, 2, 80^. The affinity of intermediate androgen metabolites are several fold lower than T and DHT^1, 2, 80^, and it is more likely that they are rapidly metabolized into T and DHT for AR binding and transcriptional activation^79, 81^. Further studies of appropriate models are needed to determine whether intermediate androgen metabolites can bind to AR and initiate its nuclear translocation in human PrCa and non-tumor cells. These studies may help identify alternative mechanisms for maintaining AR transcriptional activity that sustains PrCa growth, and result in differences in clinical outcomes in AA and EA men.

We found that the percent of AR-positive nuclei was significantly lower in Gleason score>=4+3 PrCa tissue at the time of RP in AA compared to EA men. However, the percentage of AR-positive nuclei was greater in adjacent non-tumor tissue of Gleason score>=4+3 PrCa in AA compared to EA men. In addition, DHT levels in the serum were positively correlated to AR transcriptional activity only in AA men. Therefore, our results suggest that increased AR nuclear expression in adjacent non-tumor tissue of AA men may support AR transcriptional activity for the nearby PrCa cells. There is evidence for a field effect that influences molecular changes in the tissues surrounding PrCa^82, 83^. In support of this hypothesis, previous studies observed that paracrine signaling by stromal AR promotes proliferation and invasion of PrCa cells *in vitro* ^84, 85^. This finding warrants additional studies in preclinical models of adjacent non-tumor and PrCa tissues isolated from men of different races to delineate the role of paracrine AR signaling in PrCa. Furthermore, the validation of our results in larger cohorts of primary PrCa may support the use of serum DHT levels as a surrogate marker for AR activity in treatment naïve AA men with PrCa.

To summarize, our results provide a rationale for moving treatments targeting androgen metabolizing enzymes and AR transcriptional activity earlier in the disease continuum, even in the treatment naïve setting for AA men with PrCa. In the era of personalized and precision medicine, our study along with other ongoing efforts^86–90^, emphasize the need to include underrepresented racial groups to understand and develop better prognostic and diagnostic indicators in the PrCa disease continuum.

## Methods

All the clinical samples included in Roswell Park Cohorts 1-5 were collected with informed consent at the time of radical prostatectomy at Roswell Park. Except for one patient in Roswell Park Cohort 1, all the samples were obtained prior to any treatment. The samples were de-identified before the analyses.

### RNA-sequencing, GSVA scoring and qPCR

Paired-end RNA sequencing was performed using RNA isolated from clinical samples. Pathway enrichment programs including Genego from Metacore were used to identify differentially expressed pathways between AA and EA men with PrCa. GSVA scoring was performed using the GSVA R package. GSVA scores were built on ssGSEA using a non-parametrical approach and a KS-like random walk statistic after normalizing gene expression profiles with an appropriate kernel estimation function depending on whether the expression levels were continuous or counted data. The enrichment scores compared the overall expression of two samples within the same gene-set. Given a gene-set, samples with positive enrichment values have more genes at the top the rank expression levels than samples whose genes have lower levels (for example: the bottom of the ranked expression list) with respect to the gene-set. For qPCR validation, 500ng of RNA from clinical samples were converted to cDNA using the iScript cDNA synthesis kit from Biorad (Catalog number: 1708891). Primers that overlapped intron-exon junction were designed using the NCI Primer-blast software and are listed in **Supplementary Table 13**. iTaq SYBR green kit from Biorad (Catalog number 1725121) was utilized for qPCR reactions performed on a Biorad CFX machine. The values normalized to GAPDH are presented for each primer set in PrCa and adjacent non-tumor tissues.

### Metabolism pipeline

The metabolism pipeline was performed on the Roswell Park and TCGA PrCa cohorts ^47^.

### LC-MS/MS and statistics

Study samples were analyzed for androgens (T, DHT, DHEA, ASD, AND, and 5α-dione) using a validated high pressure liquid chromatographic assay with tandem mass spectral detection (LC-MS/MS) ^49^. Immediately after the androgen analysis, the extracted samples were re-injected using a semi-quantitative targeted metabolomics approach to examine an additional 26 compounds within the steroidogenesis pathway from cholesterol to estrogens. Serum samples were quantitated using human serum calibration and quality control (QC) samples prepared by spiking known amounts of androgens into charcoal-stripped female human serum (Bioreclamation). The concentration range of the calibrators and QCs are presented in **Supplementary Table 8**. Liquid/liquid sample extractions were performed by mixing a 250 μL aliquot of a serum calibrator, QC, matrix blank, or study sample with LC/MS-grade water, internal standard (IS) solution (75.0/225 pg/mL d_3_-T/d_3_-DHT in 75% methanol), and 4.0 mL of methyl-tert-butyl ether (MTBE, EMD Millipore, Billerica, MA) in a 13 x 100 mm glass screw-top tube. Tubes were capped with teflon-lined caps, vortexed, rotated for 15 min at room temperature, and then centrifuged (Heraeus Multifuge X3R, Thermo Scientific) at 2,800 rpm and 4 °C for 15 min to separate liquid phases. The aqueous phase was frozen in a dry ice/acetone bath and the MTBE layer was poured into a clean 13 x 100 mm glass conical tube. The MTBE was evaporated at 37 °C under nitrogen and the residue was reconstituted with 60.0 μL of 60% methanol. The resulting suspensions were centrifuged (Heraeus Multifuge X3R, Thermo Scientific) at 2,800 rpm at 4 °C for 5 min to separate insoluble materials. The clear supernatant was transferred to an autosampler vial and a 7-15 μL aliquot was injected.

LC-MS/MS analyses of the extracted samples was be performed using a Prominence UFLC System (Shimadzu Scientific Instruments, Kyoto, Japan), a QTRAP® 5500 mass spectrometer (AB Sciex, Framingham, MA), with an electrospray ionization source, and two 10-port switching valves (Valco instruments Co. Inc., Houston, TX, model EPC10W). Chromatographic separation is achieved using a Phenomenex Luna C18(2) column (part number 00F-4251-B0) preceded by a Phenomenex SecurityGuard cartridge (C18, part number AJ0-4286), which were maintained at 60 °C and a flow rate of 175 µL/min. Chromatography was performed using a biphasic gradient (Mobile Phase A: 65% methanol containing 0.400 ml of 1.00 M ammonium formate and 62.0 µl of concentrated formic acid per liter; and Mobile Phase B: 100% methanol with 0.400 ml of 1.00 M ammonium formate and 62.0 µl of concentrated formic acid per liter). After the standard androgen analysis, each sample was re-injected under a different set of gradient conditions to perform semi-quantitative analysis of the steroidogenesis pathway. Analytes were detected using multiple reaction monitoring (MRM) in positive ion mode controlled by AB Sciex Analyst® software, version 1.6.2. Mass spectrometer conditions for androgens were ion spray voltage 5,250 volts, turbo gas temperature 700 °C, gas 1 = 65, gas 2 = 60, curtain gas 20, collision-associated dissociation (CAD) gas = medium. Q1 and Q3 were set at unit mass resolution, and nitrogen was used for all gases. Voltages for maximum parent/fragment ion pair intensities were optimized using direct infusion and flow injection analysis. Minor modifications in the operating conditions were performed to maintain optimal sensitivity.

Calibration curves were generated using analyte/IS peak area response ratios versus nominal concentrations (ng/mL) and weighted linear regressions using a weighting factor 1/concentration^2^. d_3_-T was used as the IS for T, ASD and DHEA, and d_3_-DHT for DHT and AND. Back-calculated concentrations were generated using the formula x = (y - b)/m, where x is the back-calculated concentration, y is the analyte/IS peak area ratio, b is the y-intercept and m is the slope. Calibrator and quality control acceptance criteria follow FDA bioanalytical guidelines whereby all acceptable concentrations must have accuracy deviations of ≤±15% from the nominal concentration with relative standard deviations (% RSD) ≤15% except at the lower limit of quantitation (LLOQ), where ≤20% deviations were allowed for both parameters. Performance data from the analysis of 855 human serum samples is provided in **Supplementary Table 9**.

The patient characteristics were summarized by race or age groups using the mean, median, and standard deviation for continuous variables; and using frequencies and relative frequencies for categorical variables (**Supplementary Tables 2, 7 and 15**). Comparisons were made using the Mann-Whitney U or Pearson chi-square tests, as appropriate. The androgen metabolites were summarized by race or age group using the mean, median, and standard deviation. To account for some observations that were BLQ, the mean and standard deviation were estimated in SAS using the maximum likelihood approach described by Croghan and Egeghy ^91^. The metabolite levels were compared using a left censored version of the log-rank test. The association between metabolite levels and other patient characteristics were evaluated using these methods. The time-to-event outcomes were summarized by race or age group using standard Kaplan-Meier methods and compared using the standard log-rank test. Overall survival (OS) was defined as the time from surgery until death due to any cause or last follow-up. Progression-free survival (PFS) was defined as the time from surgery until recurrence, death due to PrCa, or last follow-up. PFS is not calculated for patients with persistent disease. Freedom from radical prostatectomy failure (RPF) was defined as the time from surgery until persistent disease, disease progression, subsequent treatment, death from PrCa, or last follow-up. The metabolite levels were dichotomized into low (< 50th percentile) and high (> 50th percentile) with the median cut-off values, determined in the pooled samples, included in the <50 percentile group or into BLQ and detectable (i.e. >BLQ). The time-to-event outcomes were summarized for the categorized metabolite levels using standard Kaplan-Meier methods and log-rank tests. All analyses were conducted in SAS v9.4 (Cary, NC) at a significance level of 0.05.

### AIMs and SNPs

AIMs analysis was performed by Dr. Kittles group^92^. The association between AIMs and levels of androgens and intermediate metabolites in the serum were measured using Spearman correlation analysis. Best practices for germline variant calling with GATK was followed to identify the SNPs using blood whole exome data. The variants that passed the filter were kept for the association studies between race and AR metabolic genes. Fisher exact test was applied to the variants and p-values of less than 0.05 were considered statistically significant.

### Immunohistochemistry, Image J quantitation and statistics

TMA slides were stained with an AR antibody (Agilent, #M3562) and digitally scanned using Aperio Scanscope. Individual PrCa images were captured using Scanscope and the ImmunoRatio plugin (ImageJ, NIH) was used to analyze the percentage of positive AR nuclei in each clinical sample. Patient demographic and clinical characteristics for all the samples were analyzed using the mean and standard deviation for continuous variables and frequencies and relative frequencies for categorical variables. The patient demographic and clinical characteristics were compared between racial groups (EA versus AA) or age groups (<55 versus >=55) using the Mann-Whitney U and Fisher’s exact tests for continuous and categorical variables, respectively. For each patient characteristic, the AR scores and ratios were modeled as a function of the patient characteristic, race or age, and their interaction using a general linear model. F-tests about the appropriate linear combination of model estimates were used to test for: A) the association between the patient characteristic and AR within each race or age group; and B) the race or age effect on the association between AR and the patient characteristic (ie. testing the interaction term). All analyses were completed in SAS v9.4 (Cary, NC) at a significance level of 0.05. A p-value less than 0.05 denoted a statistically significant result.

## Author contributions

Conceived and designed experiments and wrote the manuscript: SR, AW; Discussed the data and revised the manuscript: SR, AW, ECG, KA, JW, SRR, DJS, GA, JLM, WJH, RAK; Performed experiments: SR, AW, RAK; Analyzed the data: SR, AW, ECG, KA, JW, SRR, GA, RAK; Contributed reagents/materials/analysis tools: SR, AW, ECG, KA, JW, SRR, DJS, JLM, WJH, RAK. All authors read and approved the final manuscript.

## Acknowledgments

We thank Dr. John Wilton, Director of the BMPK Resource ran and analyzed the serum samples for androgens and their related metabolites. We also thank Zara Kazmierczak for performing the validation of enzymes involved in androgen synthesis in clinical samples. This work was supported by: DoD Health Disparity Research Award (W81XWH-17-1-0115) to Dr. Woloszynska; Roswell Park Alliance Foundation Award to Dr. Huss. Clinical data delivery and Honest Broker services for this study were provided by the Biomedical Data Science Shared Resource. The work was also supported by the following Shared Resources: Bioanalystics, Metabolomics and Pharmacokinetics; Bioinformatics; Biostatistics; Data Bank and BioRepository Genomics and Pathology Network. The support for the Shared Resources cores are funded by the National Cancer Institute (NCI P30CA16056).

