## Supplemental files for "Racial differences in androgen metabolism and receptor signaling in prostate cancer"

#### Supplementary Figure legends

**Supplementary Figure 1 qPCR validation of RNA sequencing analysis in a subset of PrCa and adjacent non-tumor tissues from AA and EA men.** qPCR validation of RNA-sequencing results shows differential expression of enzymes involved in alternate pathways of DHT biosynthesis in a subset of PrCa and adjacent non-tumor tissues from AA and EA men.

**Supplementary Figure 2 Enzymes involved in cholesterol and androgen biosynthesis are differentially expressed in PrCa and adjacent non-tumor tissues from AA and EA men.** The heatmap represents the computed p-values obtained upon race and tissue comparisons of enzymes involved in cholesterol and androgen biosynthesis. Each column represents comparisons between tumor and adjacent non-tumor tissues or tissue-specific comparisons between AA and EA men. Dark blue rectangles represent those comparisons that are significantly different in each comparison.

**Supplementary Figure 3 Androgen biosynthesis pathways.** The illustration represents DHT and T biosynthesis from multiple pathways. The grey highlighted box represents the primary pathway of androgen biosynthesis. The blue labeled intermediate metabolites measured in the serum are significantly different in AA and EA men with PrCa.

**Supplementary Figure 4 Correlation between serum levels of androgens and intermediate metabolites.** Heatmap representing spearman correlation values between all the measured androgens and intermediate metabolites in **A.** all men, **B.** AA men, and

**C.** EA men. Asterisks in each box represent a significant correlation between the metabolites.

**Supplementary Figure 5 Overall survival and recurrence after radical prostatectomy in Roswell Park Cohort 2. A.** Overall survival and **B.** Recurrence-free survival after radical prostatectomy are not significantly different in AA and EA men with PrCa in Roswell Park Cohort 2.

**Supplementary Figure 6 Androgen receptor protein expression in PrCa from AA and EA men. A.** Representative images of AR expression in PrCa and adjacent non-tumor tissues from AA and EA men. **B.** Image J quantitation of IHC shows significantly higher AR positive nuclei in adjacent non-tumor tissues from AA compared to EA men. Each dot represents the average percent of positive AR nuclei in triplicate tissue cores. NT: Adjacent non-tumor tissue and T: PrCa tissue

**Supplementary Figure 7 Serum DHT levels are negatively correlated to AR protein expression in adjacent non-tumor and PrCa tissues only from AA men.** The top panel represents the correlation between DHT levels in the serum (Y-Axis) and the percent of positive AR nuclei (X-axis) in samples included in both Roswell Park Cohort 2 and 5. The bottom panel represents the correlation between T levels in the serum (Y-Axis) and the percent of positive AR nuclei (X-axis) in samples included in both Roswell Park Cohort 2 and 5.

**Supplementary Table Legends**

**Supplementary Table 1** Cohort names and number of patients in each cohort

**Supplementary Table 2** Patient demographics in RNA sequencing analysis

**Supplementary Table 3** Differentially expressed genes in PrCa tissues in AA and EA men from Roswell Park Cohort 1

**Supplementary Table 4** Primer sequences utilized for qPCR validation

**Supplementary Table 5** Metabolic pathways that are significantly different in PrCa and adjacent non-tumor tissues in Roswell Park Cohort 1 and TCGA

**Supplementary Table 6** Metabolic pathways that are significantly different in a race-specific manner between PrCa and adjacent non-tumor tissues in Roswell Park Cohort 1 and TCGA

**Supplementary Table 7** Patient demographics in metabolite analysis

**Supplementary Table 8** Androgen Serum calibrator QC concentrations

**Supplementary Table 9** Performance data for androgen calibrators

**Supplementary Table 10** Association of serum T and DHT levels with Gleason score

**Supplementary Table 11** Frequency of high and low levels of androgens and intermediate metabolites in the serum of AA and EA men with PrCa

**Supplementary Table 12** AIMs and self-identified race in a subset of AA and EA men with PrCa

**Supplementary Table 13** SNPs in CYP11 family of enzymes in individual samples analyzed by exome sequencing

**Supplementary Table 14** Survival rates for high vs low levels of androgen metabolites

**Supplementary Table 15** Patient demographics in TMA analysis

### Supplementary Figure 1

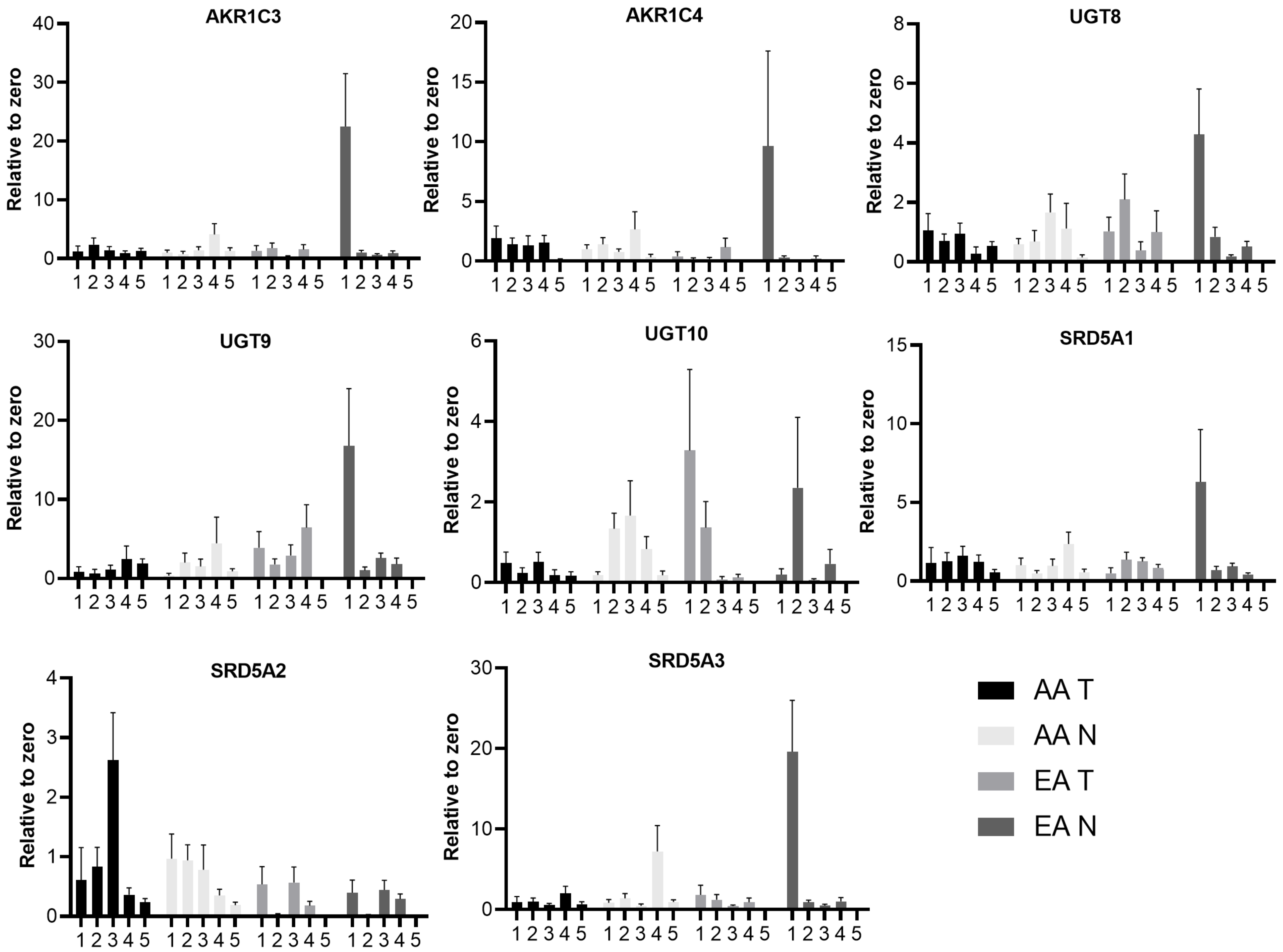

#### Supplementary Figure 2

**Steroid Synthesis**
**Race and Tumor differences**
**Pvalues**

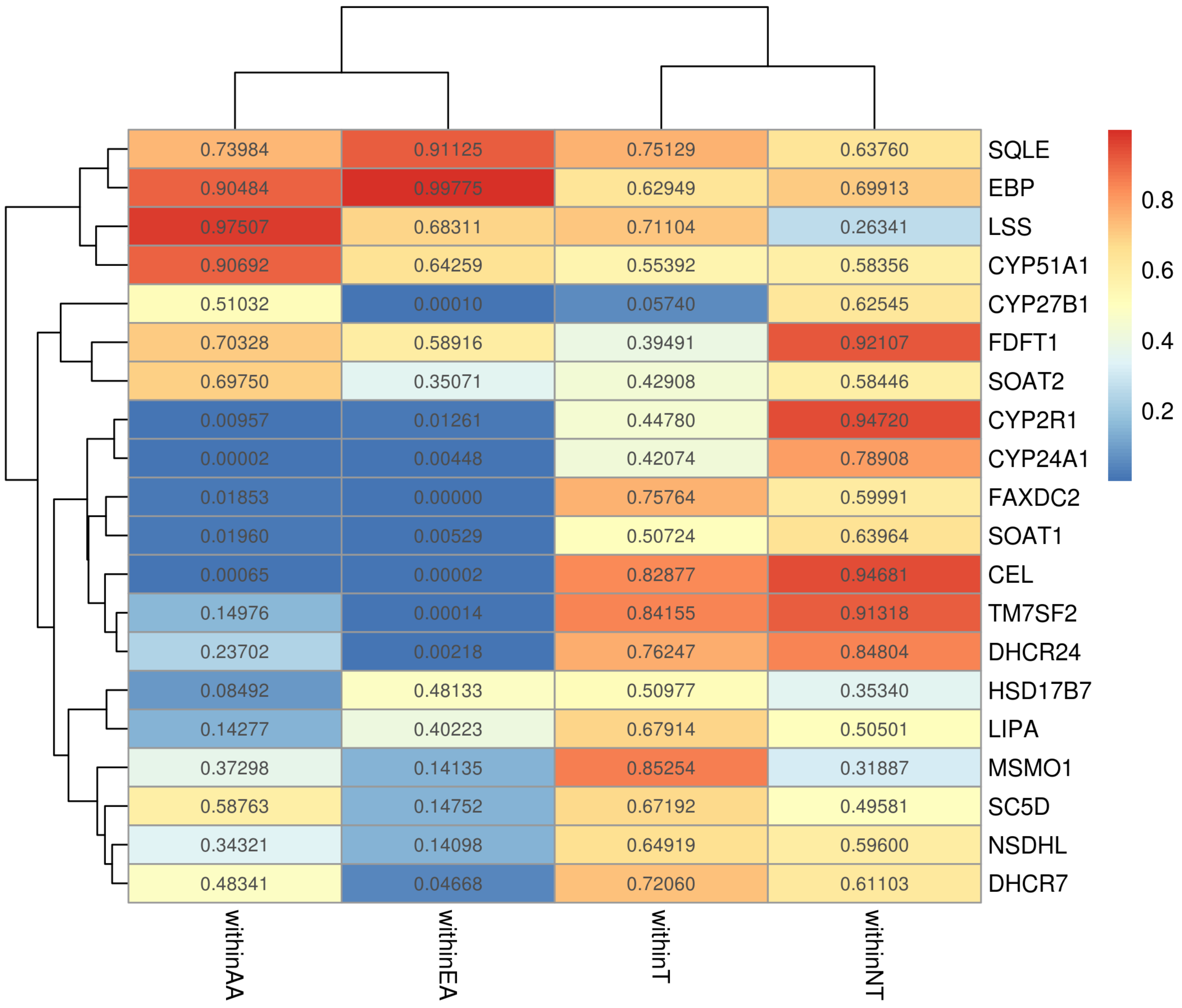

### Supplementary Figure 3

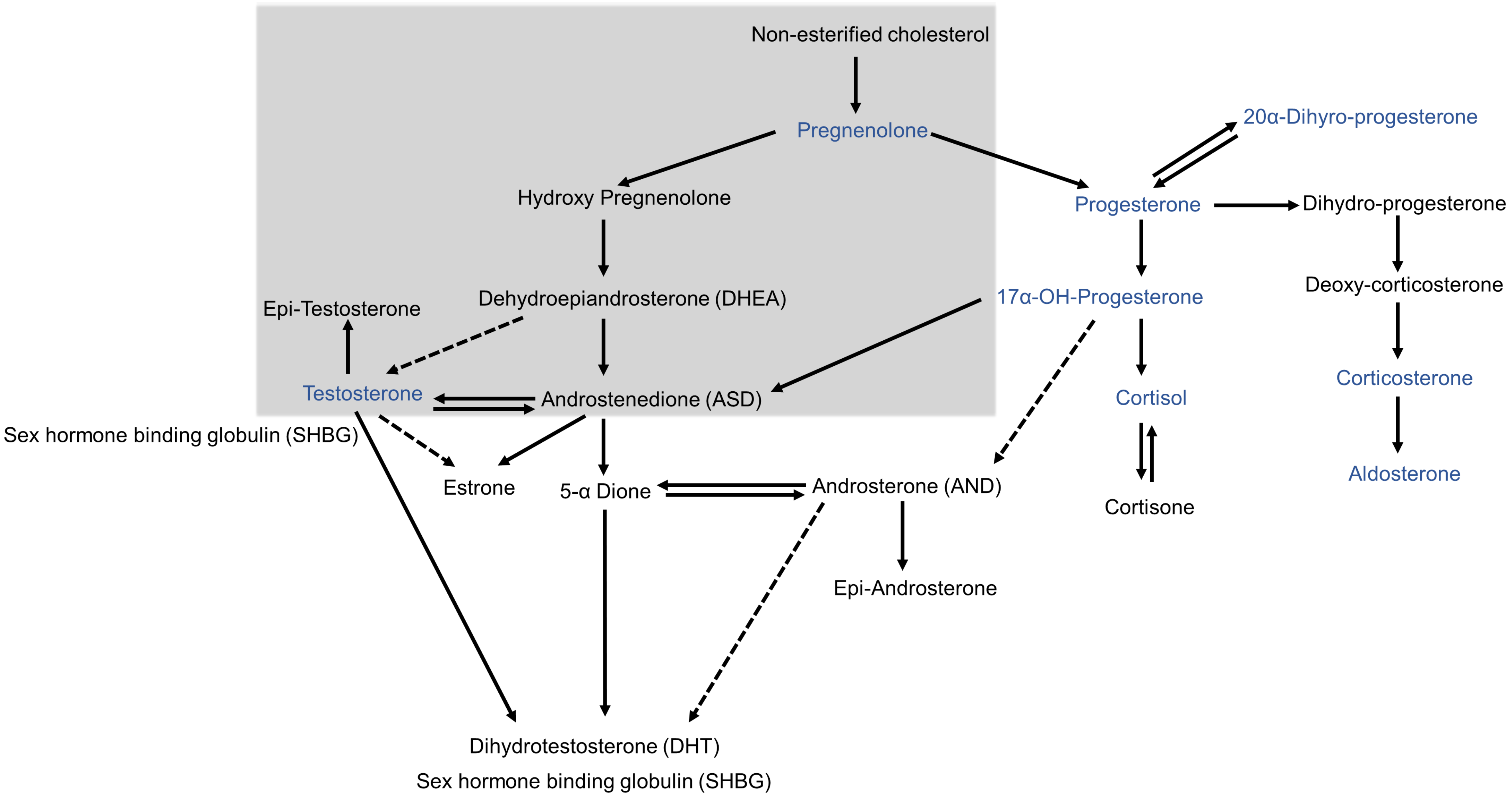

### Supplementary Figure 4

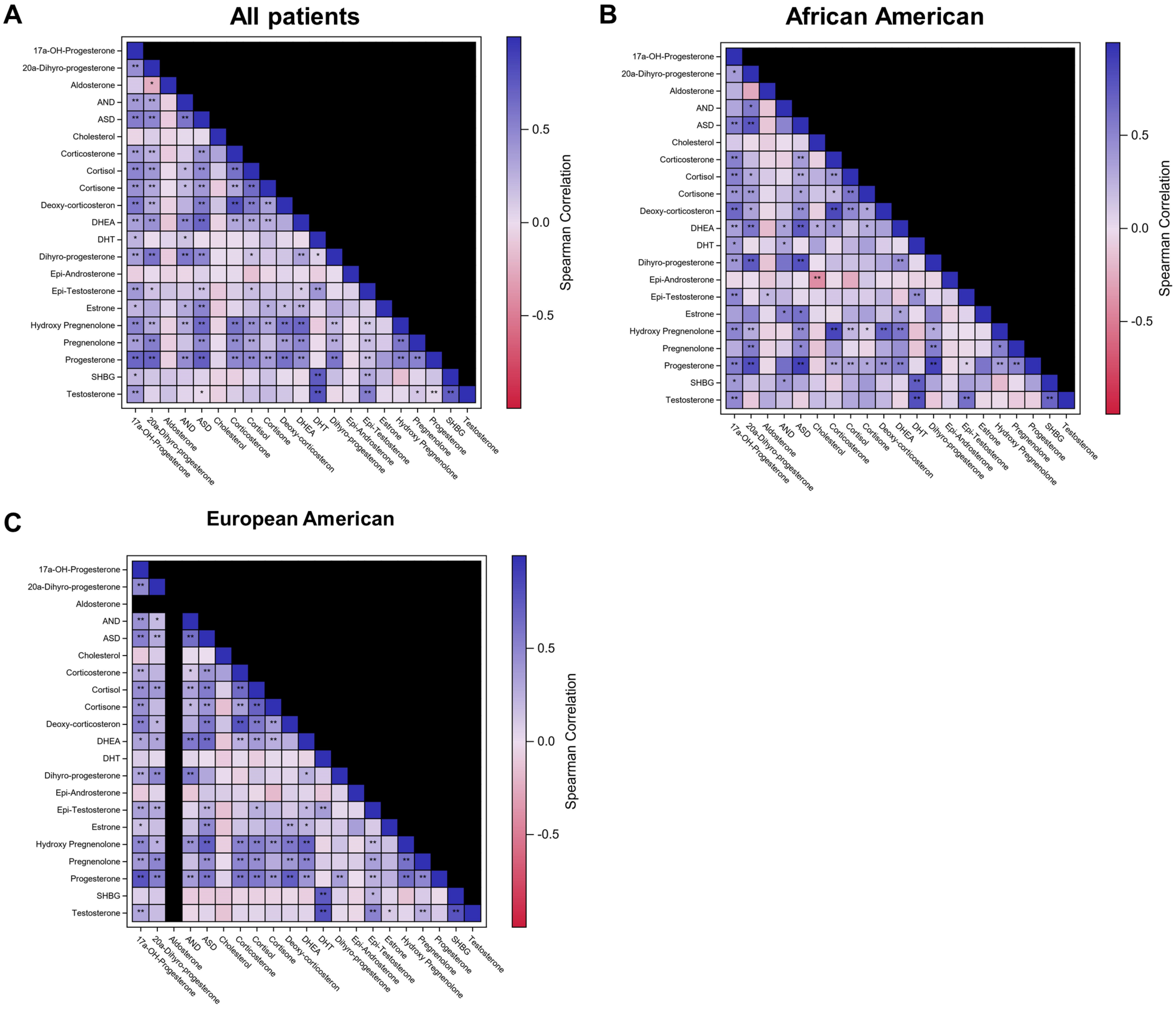

**Supplementary Figure 5**

A

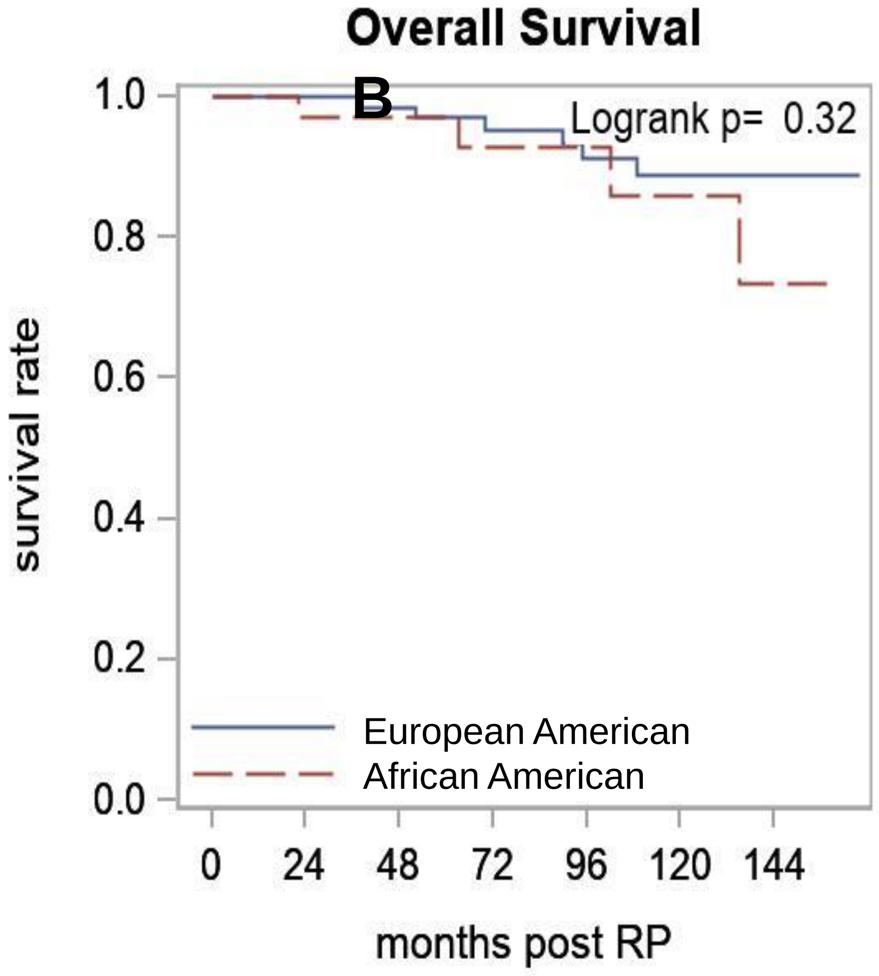

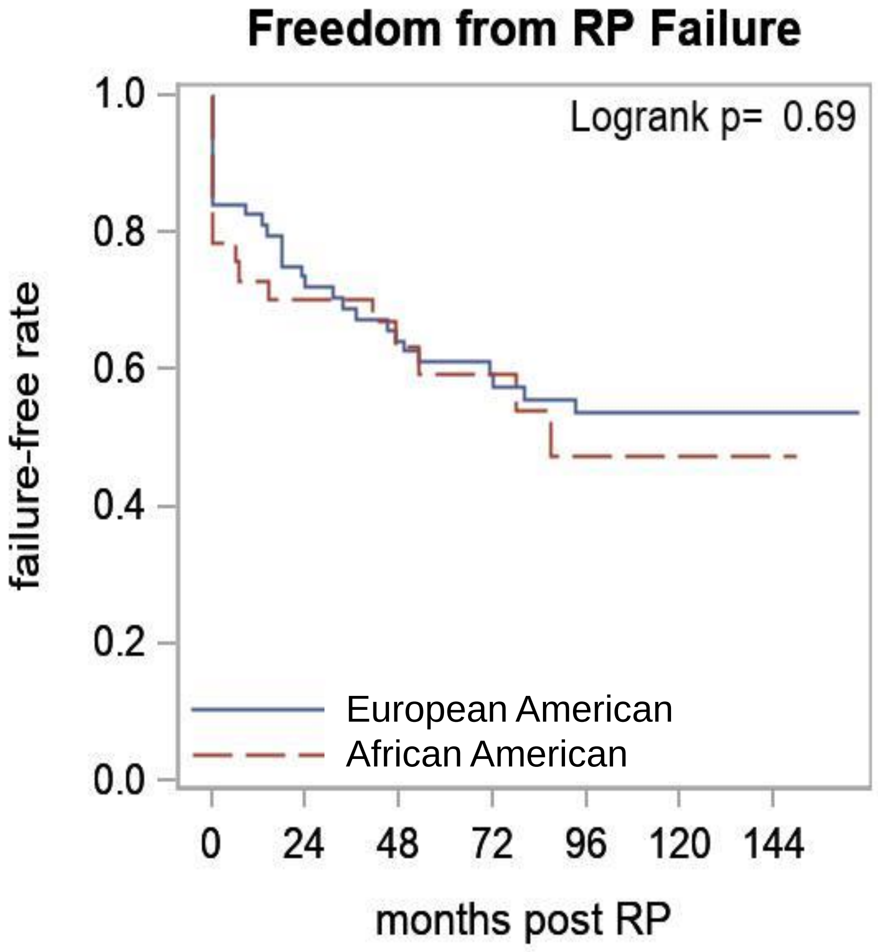

#### Supplementary Figure 6

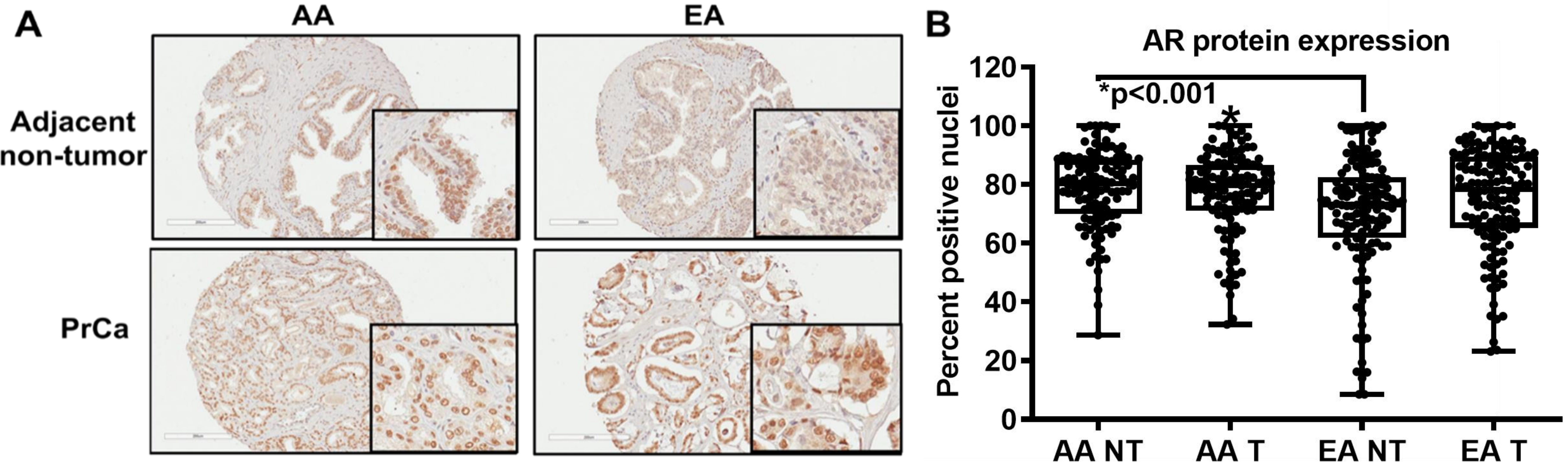

### Supplementary Figure 7

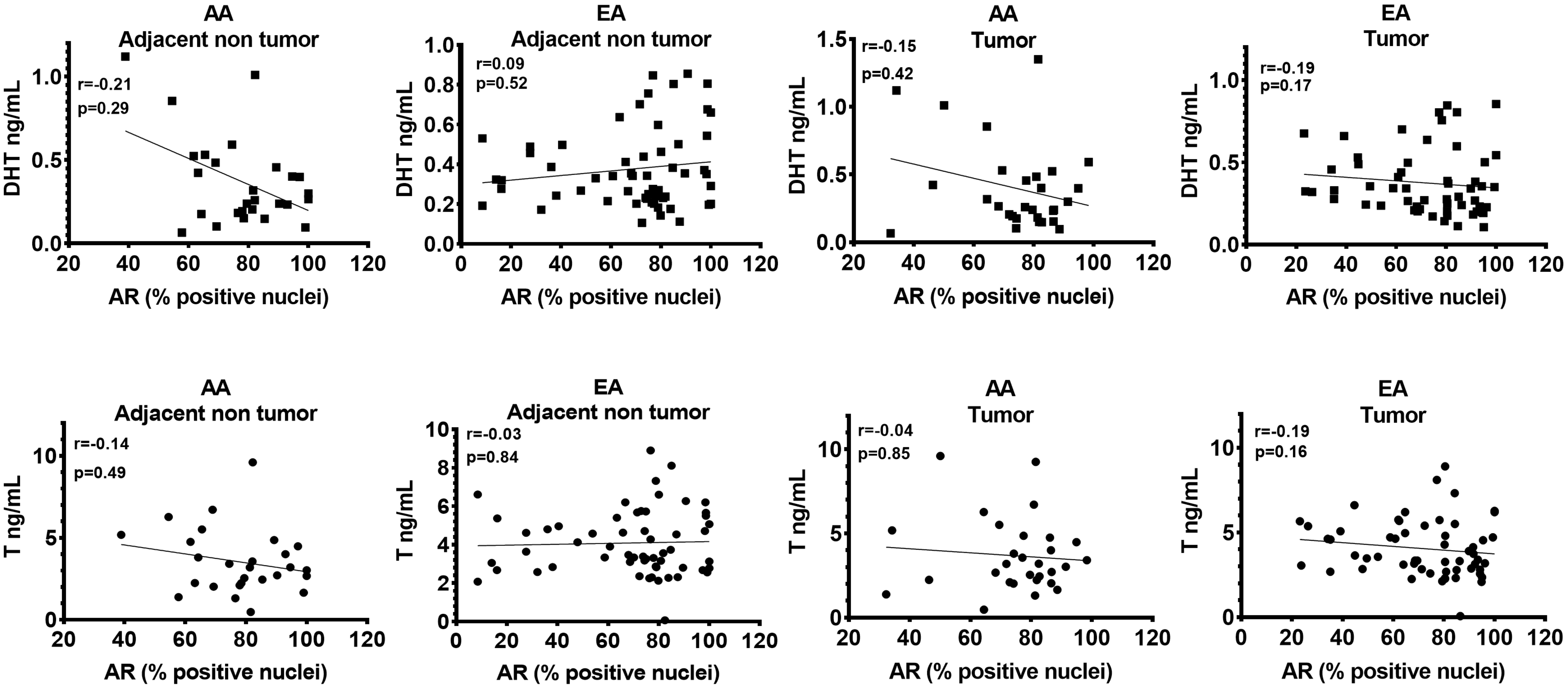

| Supplementary Table 1 Cohort names and sizes | | | |
|---|---|---|---|
| **Analysis/source** | **Cohort Name** | **AA (Total n)** | **EA (Total n)** |
| RNA sequencing | Roswell Park Cohort 1 | T=32, NT=32 | T=32, NT=32 |
| Metabolites | Roswell Park Cohort 2 | 38 | 69 |
| AIMs | Roswell Park Cohort 3 | 53 | 83 |
| Exome sequencing | Roswell Park Cohort 4 | 20 | 5 |
| TMA | Roswell Park Cohort 5 | 107 matched T and NT | 133 matched T and NT |
| The Cancer Genome Atlas | TCGA | 43 | 270 |
| **Overlapping samples (n)** | | **AA** | **EA** |
| RNA sequencing | AIMs | 6 | 5 |
| RNA sequencing | Metabolites | 8 | 11 |
| AIMs | Metabolites | 38 | 68 |
| TMA | Metabolites | 29 matched T and NT | 58 matched T and NT |
| TMA | AIMs | 41 | 69 |

| Supplementary Table 2 Demographics of patients in RNA sequencing | | | |
|---|---|---|---|
| | | **AA** | **EA** |
| Overall | N | 32 | 32 |
| Age at Diagnosis | Mean | 56.5 | 59.75 |
| | Min/Max | 44/73 | 48/72 |
| Age* | < 55 | 12 | 5 |
| | >= 55 | 20 | 27 |
| Pre-RP PSA | Mean | 10.62 | 9.08 |
| | Min/Max | 2.8/82.12 | 1.7/25.87 |
| PSA* | < 4 | 3 | 5 |
| | 4 to <10 | 24 | 17 |
| | 10 to <20 | 2 | 7 |
| | >= 20 | 3 | 3 |
| Clinical Gleason (by primary and secondary Gleason score) | <= 3+4 | 24 | 20 |
| | >=4+3 | 8 | 12 |
| Clinical Gleason (by sum) | < 7 | 8 | 9 |
| | 7 | 19 | 18 |
| | > 7 | 5 | 5 |

| | | | |
|---|---|---|---|
| | **Supplementary Table 3 Differentially expressed genes in prostate tumors fom AA and EA men (Roswell Park cohort)** | | |
| | **Upregulated pathways** | | |
| | **Name** | **p-value** | **Genes in network** |
| 1 | Protein folding and maturation POMC processing | 2.262E-23 | ACTH, PC2, alpha-MSH, proACTH, POMC, Joining peptide (JP), N-POMC, DA-alphaMSH, beta-MSH, CLIP, beta-Endorphin extracellular region, gamma-MSH, N-POC, beta-LPH, gamma-LPH, ACTH 1-17, gamma2-MSH, gamma3-MSH |
| 2 | Protein folding and maturation Posttranslational processing of neuroendocrine peptides | 2.223E-15 | PC2, CCK8-Gly, GRP(1-27), GRP(1-27)Gly, CCK8, NN, Pro-CCK, GRP precursor, NT/NN, NT, LargeNT, GRP(18-27), ProGRP, GRP(1-17), CCK8-GlyArgArg, LargeNN |
| 3 | Androstenedione and testosterone biosynthesis and metabolism p.2 | 3.500E-06 | UGT1A10, AKR1C2, AKR1C1, UGT1A9, UGT1A8, S5AR2, AKR1C4 |
| 4 | Stimulation of TGF-beta signaling in lung cancer | 6.058E-06 | TGF-beta 3, ACTA2, TGF-beta, Tropomyosin-1, SLUG, PAI1, RelA (p65 NF-kB subunit), Tropomyosin-2 |
| 5 | Cytoskeleton remodeling Keratin filaments | 8.080E-06 | Keratin 14, Keratin 16, Keratin 5/14, Keratin 4, Actin cytoskeletal, Tubulin alpha, Keratin 5 |
| 6 | Calcium-dependent regulation of normal and asthmatic smooth muscle contraction | 1.273E-05 | Endothelin-3, MYLK1, ACM2, Ryanodine receptor 2, MLCK, Neurokinin-2 receptor, MRLC, TRPC6, TRPC3 |
| 7 | Development TGF-beta-dependent induction of EMT via RhoA, PI3K and ILK | 4.334E-05 | TGF-beta 3, ACTA2, Actin, Tropomyosin-1, SLUG, RelA (p65 NF-kB subunit), MKL1 |
| 8 | Cell adhesion_Integrin-mediated cell adhesion and migration | 5.753E-05 | MyHC, MYLK1, Actin cytoskeletal, RASGRF1, MLCK, MRLC, Collagen IV |
| 9 | Development MAG-dependent inhibition of neurite outgrowth | 1.072E-04 | MyHC, TrkC, Actin cytoskeletal, TrkA, RASGRF1, MRLC |
| 10 | Airway smooth muscle contraction in asthma | 1.572E-04 | MyHC, MLCK, MRLC, TRPC6, CPI-17, TRPC3, Telokin |
| | **Downregulated pathways** | | |
| | **Name** | **p-value** | **Genes in network** |
| 1 | Development Transcription factors in segregation of hepatocytic lineage | 2.467E-06 | HNF1-beta, Activin A, p16INK4, p14ARF, OC-2, Activin |
| 2 | NETosis in SLE | 5.325E-05 | PAD4, Histone H2, Histone H2A, Histone H4, Histone H3 |
| 3 | Epigenetic alterations in ovarian cancer | 8.177E-04 | HNF1-beta, p16INK4, HRK, CDC20, SSTR1, Histone H3 |
| 4 | Development Schema: Histone H3 demethylases in stem cells | 8.487E-04 | p16INK4, p14ARF, SCN2A, Histone H3 |
| 5 | Signal transduction Activin A signaling regulation | 9.557E-04 | Activin A, Histone H2, Histone H4, Histone H3 |
| 6 | Beta-catenin-dependent transcription regulation in colorectal cancer | 1.333E-03 | Claudin-1, LAMC2, LAMC2 (80kDa), LAMC2 (100kDa) |
| 7 | Cell adhesion Tight junctions | 1.636E-03 | Claudin-8, Claudin-16, Claudin-2, Claudin-1 |
| 8 | Populations of skin dendritic cells involved in contact hypersensitivity | 1.685E-03 | CD1d, Ep-CAM, CD1a |
| 9 | Development H3K36 demethylation in stem cell maintenance | 2.308E-03 | p16INK4, p14ARF, Histone H3 |
| 10 | Development_Generation of pancreatic beta-cells from embryonic stem cells (early stages) | 2.666E-03 | HNF1-beta, Activin A, HB9 |

**Supplementary Table 4 Primer sequences**

| | Forward primer | Reverse primer |
|---|---|---|
| *AKR1C1* | TATGCGCCTGCAGAGGTTC | GGAAGCCAGCTTCAATTGCC |
| *AKR1C2* | ACCATTGGAATGACATACTGCATC | CAACCGTTTCTTACCTGTGGA |
| *AKR1C3* | AGAAGTAAAGCTTTGGAGGTCACA | AATGAGCAGAATCTATATGGCGGA |
| *AKR1C4* | TGAAAGAAGTGGCAAGCAATGG | CTGTTCCTCGGAACCTCTGG |
| *GAPDH* | GTGGTCTCCTCTGACTTCAAC | CCTGTTGCTGTAGCCAAATTC |
| *SRD5A1* | CTACGGGCATCGGTGCTTA | AATCGCCATTGTACACGCCA |
| *SRD5A2* | TGCCTTCCTTCGCGGTG | AGTACACAAATGTCCTGTGGAA |
| *SRD5A3* | TCAGGGGTTGTGAGAAAGCTG | ACCAAAGGCCATGCTACGTT |
| *UGT1A8* | TCCAAACACCTGTCACAGCA | AGGCTTCAAATTCCATAGGCA |
| *UGT1A9* | GATCTCTACAGCCACACATCA | AGGCTTCAAATTCCATAGGCA |
| *UGT1A10* | AAATTCTCCAAACCCCTGTCA | AGGCTTCAAATTCCATAGGCA |

**Supplementary Table 5 Metabolic pathways that are significantly different in prostate tumors compared to adjacent non tumor tissues**

| AA | | |
|---|---|---|
| **Roswell Park** | | |
| **Pathway** | **Pathway Scores** | **p-values** |
| Primary.Bile.Acid.Biosynthesis | 1.022263 | 0.00666 |
| Pentose.and.Glucuronate.Interconversions | 0.9147634 | 0.01979 |
| Purine.Biosynthesis | 0.7147289 | 0.02072 |
| Biotin.Metabolism | 0.20187 | 0.02636 |
| Butanoate.Metabolism | 0.6871708 | 0.03028 |
| Hexosamine.Biosynthesis | 0.3206126 | 0.03186 |
| Ascorbate.and.Aldrate.Metabolism | 0.6171322 | 0.03627 |
| Drug.Metabolism.by.other.enzymes | 0.7637301 | 0.03694 |
| Porphyrin.and.Chlorophyll.Metabolism | 0.6787258 | 0.04381 |
| Amino.Sugar.and.Nucleotide.Sugar.Metabolism | 0.6963151 | 0.04683 |
| Folate.One.Carbon.Metabolism | 0.5840967 | 0.04714 |
| **TCGA** | | |
| **Pathway** | **Pathway Scores** | **p-values** |
| Purine.Biosynthesis | 2.901336 | 0.00658 |
| Purine.Metabolism | 3.855611 | 0.03511 |
| Ascorbate.and.Aldrate.Metabolism | 1.487192 | 0.03055 |

| EA | | |
|---|---|---|
| **Roswell Park** | | |
| **Pathway** | **Pathway Scores** | **p-values** |
| N.Glycan.Biosynthesis | 4.62758 | 0.011 |
| Primary.Bile.Acid.Biosynthesis | 4.167052 | 0.00517 |
| Amino.Sugar.and.Nucleotide.Sugar.Metabolism | 3.898586 | 0.01516 |
| Folate.One.Carbon.Metabolism | 3.848161 | 0.01147 |
| Pyrimidine.Metabolism | 3.79117 | 0.02495 |
| Purine.Biosynthesis | 3.376832 | 0.00972 |
| Fatty.Acid.Degradation | 3.348506 | 0.02571 |
| Butanoate.Metabolism | 3.15107 | 0.01495 |
| Fructose.and.Mannose.Metabolism | 2.797005 | 0.02456 |
| Pentose.and.Glucuronate.Interconversions | 2.722116 | 0.02571 |
| Alanine..Aspartate.and.Glutamate.Metabolism | 2.583009 | 0.02805 |
| Valine..Leucine.and.Isoleucine.Degradation | 2.500855 | 0.03736 |
| Hexosamine.Biosynthesis | 2.436664 | 0.00757 |
| Glyoxylate.and.Dicarboxylate.Metabolism | 2.311083 | 0.03022 |
| Porphyrin.and.Chlorophyll.Metabolism | 2.101156 | 0.04494 |
| Pyrimidine.Biosynthesis | 1.73975 | 0.02441 |
| Biotin.Metabolism | 0.6464262 | 0.03729 |
| **TCGA** | | |
| **Pathway** | **Pathway Scores** | **p-values** |
| Primary.Bile.Acid.Biosynthesis | 5.935204 | 0.00768 |
| Hexosamine.Biosynthesis | 1.289549 | 0.03663 |

**Supplementary Table 6 Metabolic pathways that are significantly different in prostate tumors and adjacent non-tumor tissues from EA compared to AA men**

| Tumors | | |
|---|---|---|
| **Roswell Park** | | |
| **Pathway** | **Pathway Scores** | **p-values** |
| Purine.Metabolism | 4.08641 | 0.00266 |
| Tryptophan.Metabolism | 1.194286 | 0.01266 |
| Epinephrine.Biosynthesis | 0.6731224 | 0.00457 |
| Dopamine.Biosynthesis | 0.6730838 | 0.0032 |
| Norepinephrine.Biosynthesis | 0.6730838 | 0.00367 |
| Kynurenine.Metabolism | 0.6446137 | 0.00972 |
| Histidine.Metabolism | 0.5769484 | 0.02267 |
| Tyrosine.Metabolism | 0.512524 | 0.03997 |
| Phenylalanine.Metabolism | 0.5052332 | 0.01898 |
| **TCGA** | | |
| **Pathway** | **Pathway Scores** | **p-values** |
| Tryptophan.Metabolism | 5.576212 | 0.00002 |
| Kynurenine.Metabolism | 4.452179 | 0.00035 |
| Glycerolipid.Metabolism | 3.199096 | 0.00274 |
| Mucin.Type.O.Glycan.Biosynthesis | 1.59868 | 0.00428 |
| Glycerophospholipid.Metabolism | 1.216092 | 0.02501 |
| Retinol.Metabolism | 1.080823 | 0.02961 |
| Arachidonic.Acid.Metabolism | 1.076851 | 0.02091 |
| Linoleic.Acid.Metabolism | 1.068022 | 0.00941 |
| Ether.Lipid.Metabolism | 1.06555 | 0.01469 |
| Norepinephrine.Biosynthesis | 1.063714 | 0.00175 |
| Dopamine.Biosynthesis | 1.043327 | 0.00142 |
| Epinephrine.Biosynthesis | 1.036574 | 0.00205 |
| Tyrosine.Metabolism | 1.033785 | 0.01286 |
| alpha.Linoleic.Acid.Metabolism | 1.033018 | 0.00833 |
| Pentose.and.Glucuronate.Interconversions | 0.9391377 | 0.01477 |
| Histidine.Metabolism | 0.9190692 | 0.01049 |
| Drug.Metabolism.by.other.enzymes | 0.8579512 | 0.02356 |
| Ascorbate.and.Aldrate.Metabolism | 0.8121566 | 0.01557 |
| Steroid.Hormone.Biosynthesis | 0.781923 | 0.03414 |
| Steroid.Hormone.Metabolism | 0.766437 | 0.03779 |
| Porphyrin.and.Chlorophyll.Metabolism | 0.7650409 | 0.02618 |
| Phenylalanine.Metabolism | 0.7499468 | 0.01135 |
| Nitrogen.Metabolism | 0.6290105 | 0.01394 |
| Alanine..Aspartate.and.Glutamate.Metabolism | 0.4647017 | 0.04769 |
| Galactose.Metabolism | 0.4185877 | 0.04862 |
| Urea.Cycle | 0.2713203 | 0.03988 |

| Adjacent non-tumors | | |
|---|---|---|
| **Roswell Park** | | |
| **Pathway** | **Pathway Scores** | **p-values** |
| Estradiol.Biosynthesis | 0.247177 | 0.04084 |
| **TCGA** | | |
| **Pathway** | **Pathway Scores** | **p-values** |
| Nitrogen.Metabolism | 0.5980965 | 0.00941 |
| Arginine.and.Proline.Metabolism | 0.4501179 | 0.03954 |
| Glycogen.Degradation | 0.3835685 | 0.02153 |
| Glycogen.Biosynthesis | 0.3557748 | 0.01719 |
| Cardiolipin.Metabolism | 0.3029182 | 0.00658 |
| Retinoic.Acid.Metabolism | 0.1648934 | 0.04792 |
| Prostanoid.Biosynthesis | 0.0895125 | 0.0148 |

| Supplementary Table 7 Demographics of patients in the metabolic analysis | | | | | |
|---|---|---|---|---|---|
| | | **AA** | **EA** | **Overall** | **p-value** |
| Overall | N | 38 (35.5) | 69 (64.5) | 107 (100%) | |
| Age at Diagnosis | Mean/Std/N | 56.9/7.0/38 | 59.2/6.3/69 | 58.4/6.6/107 | 0.10 |
| | Median/Min/Max | 56.5/44.0/72.0 | 60.0/45.0/72.0 | 58.0/44.0/72.0 | |
| Age* | < 55 | 16 (42.1%) | 16 (23.2%) | 32 (29.9%) | <0.05 |
| | >= 55 | 22 (57.9%) | 53 (76.8%) | 75 (70.1%) | |
| Pre-RP PSA | Mean/Std/N | 8.3/7.7/37 | 11.1/13.7/69 | 10.1/11.9/106 | 0.36 |
| | Median/Min/Max | 6.3/0.7/35.0 | 6.4/1.3/87.9 | 6.4/0.7/87.9 | |
| PSA* | < 4 | 8 (21.6%) | 10 (14.5%) | 18 (17.0%) | 0.73 |
| | 4 to <10 | 22 (59.5%) | 40 (58.0%) | 62 (58.5%) | |
| | 10 to <20 | 4 (10.8%) | 11 (15.9%) | 15 (14.2%) | |
| | >= 20 | 3 (8.1%) | 8 (11.6%) | 11 (10.4%) | |
| Clinical Gleason (by primary and secondary Gleason score) | <= 3+4 | 16 (61.5%) | 27 (71.1%) | 43 (67.2%) | 0.59 |
| | >=4+3 | 10 (38.5%) | 11 (28.9%) | 21 (32.8%) | |
| Clinical Gleason (by sum) | < 7 | 9 (34.6%) | 19 (50.0%) | 28 (43.8%) | 0.28 |
| | 7 | 10 (38.5%) | 14 (36.8%) | 24 (37.5%) | |
| | > 7 | 7 (26.9%) | 5 (13.2%) | 12 (18.8%) | |
| Post-RP Hormone | No | 36 (94.7%) | 53 (76.8%) | 89 (83.2%) | <0.05 |
| | Yes | 2 (5.3%) | 16 (23.2%) | 18 (16.8%) | |

| Supplementary Table 8 Androgen Serum Calibrator QC Concentrations | | | |
|---|---|---|---|
| **Compound** | **Serum Calibration Ranges** | **Lower Limit of Quantitation** | **QC Concentrations (ng/mL)** |
| **ASD** | 0.00625 - 3.75 ng/mL | 0.00625 ng/mL | 0.0280, 0.280, 2.80 |
| **T** | 0.00625 - 3.75 ng/mL | 0.00625 ng/mL | 0.0280, 0.280, 2.80 |
| **DHEA** | 0.200 - 7.50 ng/mL | 0.200 ng/mL | 0.560, 1.68, 5.60 |
| **DHT** | 0.0125 - 7.50 ng/mL | 0.0125 ng/mL | 0.560, 0.560, 5.60 |
| **AND** | 0.200 - 7.50 ng/mL | 0.200 ng/mL | 0.560, 1.68, 5.60 |
| **5α-dione[1]** | 0.200 - 7.50 ng/mL | 0.200 ng/mL | 0.560, 1.68, 5.60 |

[1]5α-dione did not pass standard acceptance criteria during assay validation (i.e., the theoretical concentration ± 15% as recommended by FDA's Bioanalytical Guidance)

**Supplementary Table 9**

| Mass Spectral Analyte | | |
|---|---|---|
| **Analyte** | **Q1** | **Q3** |
| ASD | 287.2 | 97.1 |
| T | 289.2 | 97 |
| DHEA-1 | 289.1 | 271.2 |
| 5α-dione-1 | 289.4 | 271.2 |
| DHT | 291.1 | 255.2 |
| AND-1 | 291.4 | 273.3 |
| DHEA-2* | 289.2 | 253.2 |
| 5α-dione-2* | 289.2 | 253.2 |
| AND-2* | 291.2 | 255.3 |
| $d_3$-T | 292.2 | 97 |
| $d_3$-DHT | 294.2 | 258.2 |

*Confirmatory transitions

| Performance Data for Androgen Calibrators and from the Analysis of 855 human Serum Samples | | | | |
|---|---|---|---|---|
| **Compound** | **Calibrator Accuracy (%)** | | **Calibrator Precision (%)** | |
| | **Mean** | **Range** | **Mean** | **Range** |
| **ASD** | 100 | 95.1-103 | 2.47 | 1.86-3.23 |
| **T** | 100 | 93.2-103 | 2.08 | 1.37-3.30 |
| **DHEA** | 100 | 99.0-101 | 2.85 | 1.94-4.85 |
| **DHT** | 100 | 95.4-103 | 3.03 | 1.68-4.90 |
| **AND** | 100 | 97.0-104 | 3.96 | 2.26-5.09 |
| **5α-dione**[1] | 100 | 97.7-102 | 3.89 | 2.62-5.48 |

| **Compound** | **QC Accuracy (%)** | | **QC Precision (%)** | |
|---|---|---|---|---|
| | Mean | Range | Mean | Range |
| **ASD** | 102 | 98.6-105 | 5.72 | 4.76-6.87 |
| **T** | 105 | 99.4-108 | 4.37 | 2.95-5.49 |
| **DHEA** | 102 | 101-102 | 4.31 | 3.73-4.99 |
| **DHT** | 104 | 100-107 | 5.37 | 4.60-5.92 |
| **AND** | 101 | 97.9-103 | 7.87 | 7.73-8.02 |
| **5α-dione**[1] | 103 | 102-103 | 11.7 | 8.32-14.4 |

| Supplementary Table 10 Association of serum T and DHT levels with Gleason score | | | | |
|---|---|---|---|---|
| | Gleason Score | | | |
| **Metabolite** | **< 7 (Median/Min/Max)** | **7 (Median/Min/Max)** | **> 7 (Median/Min/Max)** | **p-value** |
| Dihydrotestosterone (DHT) (ng/mL)**^^** | 0.40 (BLQ-1.01) | 0.22 (0.07-0.70) | 0.25 (0.10-0.46) | 0.053 |
| Testosterone (T) (ng/mL)**^^** | 4.67 (0.07-9.61) | 2.50 (0.48-6.61) | 2.72 (1.65-4.86) | <0.01 |

**Supplementary Table 11 Frequency of high and low levels of androgens and related metabolites in the serum of AA and EA men with PrCa**

| | | AA | EA | Overall | p-value |
|---|---|---|---|---|---|
| 17α-OH-Progesterone (ng/mL) | Low | 26 (68.4%) | 28 (40.6%) | 54 (50.5%) | <0.01 |
| | High | 12 (31.6%) | 41 (59.4%) | 53 (49.5%) | |
| 20α-Dihyro-progesterone (ng/mL) | Low | 24 (63.2%) | 30 (43.5%) | 54 (50.5%) | 0.07 |
| | High | 14 (36.8%) | 39 (56.5%) | 53 (49.5%) | |
| Aldosterone (ng/mL) | Low | 34 (89.5%) | 69 (100.0%) | 103 (96.3%) | <0.05 |
| | High | 4 (10.5%) | | 4 (3.7%) | |
| AND* | Low | 20 (52.6%) | 34 (49.3%) | 54 (50.5%) | 0.84 |
| | High | 18 (47.4%) | 35 (50.7%) | 53 (49.5%) | |
| ASD (ng/mL) | Low | 25 (65.8%) | 29 (42.0%) | 54 (50.5%) | <0.05 |
| | High | 13 (34.2%) | 40 (58.0%) | 53 (49.5%) | |
| Corticosterone (ng/mL) | Low | 28 (73.7%) | 26 (37.7%) | 54 (50.5%) | <.001 |
| | High | 10 (26.3%) | 43 (62.3%) | 53 (49.5%) | |
| Cortisol (ng/mL) | Low | 27 (71.1%) | 27 (39.1%) | 54 (50.5%) | <0.01 |
| | High | 11 (28.9%) | 42 (60.9%) | 53 (49.5%) | |
| Cortisone (ng/mL) | Low | 22 (57.9%) | 32 (46.4%) | 54 (50.5%) | 0.31 |
| | High | 16 (42.1%) | 37 (53.6%) | 53 (49.5%) | |
| DHEA (ng/mL) | Low | 24 (63.2%) | 30 (44.1%) | 54 (50.9%) | 0.07 |
| | High | 14 (36.8%) | 38 (55.9%) | 52 (49.1%) | |
| Deoxy-corticosterone (ng/mL) | Low | 24 (63.2%) | 30 (43.5%) | 54 (50.5%) | 0.07 |
| | High | 14 (36.8%) | 39 (56.5%) | 53 (49.5%) | |
| Dihyro-progesterone (ng/mL) | Low | 21 (55.3%) | 33 (47.8%) | 54 (50.5%) | 0.55 |
| | High | 17 (44.7%) | 36 (52.2%) | 53 (49.5%) | |
| DHT (ng/mL) | Low | 22 (57.9%) | 32 (46.4%) | 54 (50.5%) | 0.31 |
| | High | 16 (42.1%) | 37 (53.6%) | 53 (49.5%) | |
| epi-Androsterone (ng/mL) | Low | 18 (50.0%) | 34 (50.7%) | 52 (50.5%) | 1.00 |
| | High | 18 (50.0%) | 33 (49.3%) | 51 (49.5%) | |
| epi-Testosterone (ng/mL) | Low | 21 (55.3%) | 33 (47.8%) | 54 (50.5%) | 0.55 |
| | High | 17 (44.7%) | 36 (52.2%) | 53 (49.5%) | |
| Estrone (ng/mL) | Low | 20 (52.6%) | 36 (52.2%) | 56 (52.3%) | 1.00 |
| | High | 18 (47.4%) | 33 (47.8%) | 51 (47.7%) | |
| Hydroxy Pregnenolone (ng/mL) | Low | 24 (63.2%) | 30 (43.5%) | 54 (50.5%) | 0.07 |
| | High | 14 (36.8%) | 39 (56.5%) | 53 (49.5%) | |
| Non-esterified cholesterol (µg/mL) | Low | 21 (55.3%) | 33 (47.8%) | 54 (50.5%) | 0.55 |
| | High | 17 (44.7%) | 36 (52.2%) | 53 (49.5%) | |
| Pregnenolone (ng/mL) | Low | 25 (65.8%) | 29 (42.0%) | 54 (50.5%) | <0.05 |
| | High | 13 (34.2%) | 40 (58.0%) | 53 (49.5%) | |
| Progesterone (ng/mL) | Low | 27 (71.1%) | 27 (39.1%) | 54 (50.5%) | <0.01 |
| | High | 11 (28.9%) | 42 (60.9%) | 53 (49.5%) | |
| SHBG (nmol/L) | Low | 20 (52.6%) | 34 (49.3%) | 54 (50.5%) | 0.84 |
| | High | 18 (47.4%) | 35 (50.7%) | 53 (49.5%) | |
| T (ng/mL) | Low | 20 (52.6%) | 34 (49.3%) | 54 (50.5%) | 0.84 |
| | High | 18 (47.4%) | 35 (50.7%) | 53 (49.5%) | |

**Supplementary Table 12 Ancestry Informative Markers and Self-Identified Race in a subset of patient population**

| WAF | NAT | EURO | Self-identified race |
|---|---|---|---|
| 0.9726 | 0.0129 | 0.0145 | African American |
| 0.97 | 0.02 | 0.01 | African American |
| 0.957 | 0.0049 | 0.038 | African American |
| 0.94 | 0.04 | 0.02 | African American |
| 0.9238 | 0.0279 | 0.0483 | African American |
| 0.91 | 0.01 | 0.08 | African American |
| 0.9095 | 0.0412 | 0.0493 | African American |
| 0.91 | 0.06 | 0.03 | African American |
| 0.8956 | 0.0776 | 0.0267 | African American |
| 0.8956 | 0.0776 | 0.0267 | African American |
| 0.88 | 0.01 | 0.10 | African American |
| 0.861 | 0.0184 | 0.1207 | African American |
| 0.8589 | 0.0272 | 0.1139 | African American |
| 0.8492 | 0.1412 | 0.0096 | African American |
| 0.85 | 0.02 | 0.13 | African American |
| 0.8442 | 0.016 | 0.1398 | African American |
| 0.8293 | 0.0452 | 0.1255 | African American |
| 0.83 | 0.03 | 0.14 | African American |
| 0.8158 | 0.0145 | 0.1697 | African American |
| 0.8156 | 0.0362 | 0.1483 | African American |
| 0.81 | 0.02 | 0.17 | African American |
| 0.8015 | 0.0064 | 0.1921 | African American |
| 0.7993 | 0.0192 | 0.1814 | African American |
| 0.79 | 0.0042 | 0.2058 | African American |
| 0.7796 | 0.0079 | 0.2125 | African American |
| 0.77 | 0.02 | 0.21 | African American |
| 0.7561 | 0.0199 | 0.224 | African American |
| 0.76 | 0.02 | 0.23 | African American |
| 0.7319 | 0.0093 | 0.2589 | African American |
| 0.73 | 0.01 | 0.26 | African American |
| 0.7295 | 0.0049 | 0.2657 | African American |
| 0.72 | 0.01 | 0.27 | African American |
| 0.68 | 0.03 | 0.29 | African American |
| 0.6573 | 0.0092 | 0.3336 | African American |
| 0.6062 | 0.1598 | 0.234 | African American |
| 0.53 | 0.01 | 0.46 | African American |
| 0.1782 | 0.0073 | 0.8145 | European American |
| 0.08 | 0.02 | 0.90 | European American |
| 0.07 | 0.02 | 0.91 | European American |
| 0.0658 | 0.0033 | 0.9309 | European American |
| 0.0636 | 0.0076 | 0.9289 | European American |
| 0.06 | 0.01 | 0.92 | European American |

| | | | |
|---|---|---|---|
| 0.06 | 0.04 | 0.90 | European American |
| 0.0601 | 0.0058 | 0.934 | European American |
| 0.0509 | 0.0219 | 0.9272 | European American |
| 0.0494 | 0.0135 | 0.9371 | European American |
| 0.0469 | 0.0053 | 0.9478 | European American |
| 0.04 | 0.01 | 0.95 | European American |
| 0.0398 | 0.013 | 0.9472 | European American |
| 0.031 | 0.0026 | 0.9664 | European American |
| 0.03 | 0.0516 | 0.9184 | European American |
| 0.0299 | 0.0251 | 0.945 | European American |
| 0.0298 | 0.0419 | 0.9282 | European American |
| 0.0289 | 0.0117 | 0.9595 | European American |
| 0.0286 | 0.0263 | 0.9451 | European American |
| 0.0275 | 0.0071 | 0.9654 | European American |
| 0.0275 | 0.0084 | 0.9641 | European American |
| 0.0245 | 0.0257 | 0.9498 | European American |
| 0.0241 | 0.0199 | 0.9559 | European American |
| 0.0224 | 0.0075 | 0.97 | European American |
| 0.0202 | 0.0852 | 0.8946 | European American |
| 0.02 | 0.05 | 0.93 | European American |
| 0.0182 | 0.0099 | 0.9718 | European American |
| 0.0162 | 0.138 | 0.8458 | European American |
| 0.02 | 0.05 | 0.93 | European American |
| 0.02 | 0.01 | 0.97 | European American |
| 0.0152 | 0.0018 | 0.983 | European American |
| 0.0149 | 0.0102 | 0.975 | European American |
| 0.0143 | 0.006 | 0.9797 | European American |
| 0.0131 | 0.0184 | 0.9685 | European American |
| 0.0128 | 0.0256 | 0.9615 | European American |
| 0.01 | 0.07 | 0.92 | European American |
| 0.0114 | 0.0254 | 0.9632 | European American |
| 0.01 | 0.06 | 0.93 | European American |
| 0.0102 | 0.0098 | 0.98 | European American |
| 0.0089 | 0.0365 | 0.9546 | European American |
| 0.01 | 0.12 | 0.87 | European American |
| 0.0082 | 0.0231 | 0.9688 | European American |
| 0.01 | 0.01 | 0.98 | European American |
| 0.0074 | 0.0281 | 0.9646 | European American |
| 0.007 | 0.0092 | 0.9837 | European American |
| 0.01 | 0.04 | 0.95 | European American |
| 0.0053 | 0.0586 | 0.9361 | European American |
| 0.005 | 0.0655 | 0.9294 | European American |
| 0.0047 | 0.0225 | 0.9727 | European American |
| 0.0046 | 0.0885 | 0.9069 | European American |
| 0.0046 | 0.0051 | 0.9903 | European American |
| 0.0046 | 0.0051 | 0.9903 | European American |
| 0.004 | 0.1154 | 0.8805 | European American |

| | | | |
|---|---|---|---|
| 0.0039 | 0.1436 | 0.8525 | European American |
| 0.0038 | 0.0654 | 0.9308 | European American |
| 0.0026 | 0.0027 | 0.9947 | European American |
| 0.0022 | 0.0161 | 0.9817 | European American |
| 0.0021 | 0.1135 | 0.8843 | European American |
| 0.0018 | 0.183 | 0.8152 | European American |
| 0.0018 | 0.0052 | 0.993 | European American |
| 0.00 | 0.01 | 0.99 | European American |
| 0.0015 | 0.0021 | 0.9964 | European American |
| 0.0009 | 0.0023 | 0.9967 | European American |
| 0.0008 | 0.0013 | 0.9979 | European American |
| 0.0008 | 0.0052 | 0.9939 | European American |

Supplementary Table 13 CYP11B family SNPs

| SNP ID | Chr | Start | End | Ref | Alt | Func.refGene | Gene.refGene |
|---|---|---|---|---|---|---|---|
| rs113759408 | 8 | 143958342 | 143958342 | G | A | intronic | CYP11B1 |
| | **0/0** | **0/1** | **1/1** | | | **Alternate** | **Wild type** |
| **AA** | 7 | 8 | 3 | | **AA** | 11 | 7 |
| **EA** | 5 | 0 | 0 | | **EA** | 0 | 5 |
| **p-value** | 0.06 | | | | **p-value** | <0.05 | |

| SNP ID | Chr | Start | End | Ref | Alt | Func.refGene | Gene.refGene |
|---|---|---|---|---|---|---|---|
| rs7818953 | 8 | 143959289 | 143959289 | T | C | intronic | CYP11B1 |
| | **0/0** | **0/1** | **1/1** | | | | |
| **AA** | 0 | 2 | 16 | | | | |
| **EA** | 1 | 2 | 2 | | | | |
| **p-value** | <0.05 | | | | | | |

| SNP ID | Chr | Start | End | Ref | Alt | Func.refGene | Gene.refGene |
|---|---|---|---|---|---|---|---|
| rs5313 | 8 | 143994253 | 143994253 | C | T | exonic | CYP11B2 |
| | **0/0** | **0/1** | **1/1** | | | **Alternate** | **Wild type** |
| **AA** | 6 | 8 | 4 | | **AA** | 12 | 6 |
| **EA** | 5 | 0 | 0 | | **EA** | 0 | 5 |
| **p-value** | 0.05 | | | | **p-value** | <0.05 | |

| SNP ID | Chr | Start | End | Ref | Alt | Func.refGene | Gene.refGene |
|---|---|---|---|---|---|---|---|
| rs4544 | 8 | 143994806 | 143994806 | A | G | exonic | CYP11B2 |
| | **0/0** | **0/1** | **1/1** | | | **Alternate** | **Wild type** |
| **AA** | 6 | 7 | 5 | | **AA** | 12 | 6 |
| **EA** | 5 | 0 | 0 | | **EA** | 0 | 5 |
| **p-value** | 0.05 | | | | **p-value** | <0.05 | |

| SNP ID | Chr | Start | End | Ref | Alt | Func.refGene | Gene.refGene |
|---|---|---|---|---|---|---|---|
| rs57166526 | 8 | 143995880 | 143995880 | G | A | intronic | CYP11B2 |
| | **0/0** | **0/1** | **1/1** | | | **Alternate** | **Wild type** |
| **AA** | 6 | 8 | 4 | | **AA** | 12 | 6 |
| **EA** | 5 | 0 | 0 | | **EA** | 0 | 5 |
| **p-value** | 0.05 | | | | **p-value** | <0.05 | |

| | |
|---|---|
| 0/0 | Wild type |
| 0/1 | One allele |
| 1/1/ | Both alleles |

| Supplementary Table 14 Survival rates for high vs low levels of androgen metabolites | | | | | | | | | | | |
|---|---|---|---|---|---|---|---|---|---|---|---|
| | | PFS | | | | | RPF | | | | |
| | | 3 years | | 5 years | | Sample size | 3 years | | 5 years | | Sample size |
| | | Rate (95% CI) | p-value | Rate (95% CI) | p-value | | Rate (95% CI) | p-value | Rate (95% CI) | p-value | |
| 17α-OH-Progesterone (ng/mL) | High | 0.96 (0.84, 0.99) | 0.07 | 0.84 (0.70, 0.92) | 0.23 | E=10 C=38 T=48 | 0.81 (0.67, 0.89) | <0.01 | 0.70 (0.56, 0.81) | <0.05 | E=19 C=34 T=53 |
| | Low | 0.84 (0.68, 0.92) | | 0.74 (0.56, 0.86) | | E=11 C=29 T=40 | 0.58 (0.43, 0.70) | | 0.51 (0.36, 0.64) | | E=27 C=27 T=54 |
| 20α-Dihyro-progesterone (ng/mL) | High | 0.88 (0.74, 0.95) | 0.44 | 0.75 (0.58, 0.86) | 0.23 | E=13 C=30 T=43 | 0.68 (0.53, 0.78) | 0.67 | 0.57 (0.42, 0.69) | 0.43 | E=26 C=27 T=53 |
| | Low | 0.93 (0.80, 0.98) | | 0.85 (0.69, 0.93) | | E=8 C=37 T=45 | 0.71 (0.57, 0.82) | | 0.65 (0.50, 0.76) | | E=20 C=34 T=54 |
| Aldosterone (ng/mL) | <BLQ | 0.90 (0.81, 0.95) | 0.53 | 0.79 (0.68, 0.87) | 0.40 | E=21 C=63 T=84 | 0.68 (0.58, 0.76) | 0.22 | 0.59 (0.49, 0.68) | 0.17 | E=46 C=57 T=103 |
| | >BLQ | 1.00 (1.00, 1.00) | | 1.00 (1.00, 1.00) | | E=0 C=4 T=4 | 1.00 (1.00, 1.00) | | 1.00 (1.00, 1.00) | | E=0 C=4 T=4 |
| AND (ng/mL) | High | 0.90 (0.76, 0.96) | 0.90 | 0.76 (0.59, 0.87) | 0.47 | E=12 C=30 T=42 | 0.68 (0.53, 0.78) | 0.63 | 0.57 (0.42, 0.69) | 0.41 | E=25 C=28 T=53 |
| | Low | 0.91 (0.77, 0.96) | | 0.83 (0.68, 0.92) | | E=9 C=37 T=46 | 0.71 (0.57, 0.82) | | 0.65 (0.50, 0.76) | | E=21 C=33 T=54 |
| ASD (ng/mL) | High | 0.94 (0.81, 0.98) | 0.32 | 0.81 (0.66, 0.90) | 0.69 | E=11 C=37 T=48 | 0.79 (0.65, 0.88) | <0.05 | 0.65 (0.50, 0.77) | 0.19 | E=21 C=32 T=53 |
| | Low | 0.87 (0.72, 0.94) | | 0.78 (0.61, 0.89) | | E=10 C=30 T=40 | 0.60 (0.46, 0.72) | | 0.56 (0.41, 0.68) | | E=25 C=29 T=54 |
| Corticosterone (ng/mL) | High | 0.93 (0.80, 0.98) | 0.37 | 0.91 (0.77, 0.96) | <0.05 | E=6 C=40 T=46 | 0.73 (0.58, 0.83) | 0.38 | 0.68 (0.54, 0.79) | 0.13 | E=19 C=34 T=53 |
| | Low | 0.88 (0.73, 0.95) | | 0.68 (0.50, 0.80) | | E=15 C=27 T=42 | 0.66 (0.51, 0.77) | | 0.53 (0.38, 0.66) | | E=27 C=27 T=54 |
| Cortisol (ng/mL) | High | 0.89 (0.76, 0.95) | 0.65 | 0.80 (0.65, 0.89) | 0.92 | E=13 C=35 T=48 | 0.75 (0.61, 0.85) | 0.18 | 0.66 (0.52, 0.78) | 0.18 | E=22 C=31 T=53 |
| | Low | 0.92 (0.77, 0.97) | | 0.80 (0.62, 0.90) | | E=8 C=32 T=40 | 0.64 (0.49, 0.75) | | 0.55 (0.40, 0.67) | | E=24 C=30 T=54 |
| Cortisone (ng/mL) | High | 0.89 (0.76, 0.95) | 0.57 | 0.80 (0.65, 0.89) | 0.78 | E=12 C=36 T=48 | 0.74 (0.60, 0.84) | 0.24 | 0.66 (0.51, 0.77) | 0.26 | E=20 C=33 T=53 |
| | Low | 0.92 (0.78, 0.97) | | 0.80 (0.63, 0.90) | | E=9 C=31 T=40 | 0.65 (0.50, 0.76) | | 0.56 (0.41, 0.68) | | E=26 C=28 T=54 |
| DHEA (ng/mL) | High | 0.93 (0.80, 0.98) | 0.59 | 0.81 (0.65, 0.90) | 0.94 | E=10 C=35 T=45 | 0.75 (0.60, 0.84) | 0.29 | 0.62 (0.47, 0.74) | 0.77 | E=21 C=31 T=52 |
| | Low | 0.90 (0.75, 0.96) | | 0.81 (0.64, 0.90) | | E=10 C=32 T=42 | 0.66 (0.51, 0.77) | | 0.61 (0.46, 0.73) | | E=24 C=30 T=54 |
| DHT (ng/mL) | High | 0.89 (0.75, 0.95) | 0.50 | 0.81 (0.65, 0.90) | 0.96 | E=12 C=32 T=44 | 0.73 (0.59, 0.83) | 0.40 | 0.67 (0.52, 0.78) | 0.23 | E=22 C=31 T=53 |
| | Low | 0.93 (0.79, 0.98) | | 0.79 (0.63, 0.89) | | E=9 C=35 T=44 | 0.65 (0.51, 0.77) | | 0.54 (0.39, 0.67) | | E=24 C=30 T=54 |
| Deoxy-corticosterone (ng/mL) | High | 0.93 (0.80, 0.98) | 0.39 | 0.91 (0.77, 0.96) | <0.05 | E=6 C=40 T=46 | 0.73 (0.58, 0.83) | 0.40 | 0.71 (0.56, 0.81) | 0.07 | E=18 C=35 T=53 |
| | Low | 0.88 (0.73, 0.95) | | 0.68 (0.50, 0.80) | | E=15 C=27 T=42 | 0.66 (0.51, 0.77) | | 0.51 (0.36, 0.64) | | E=28 C=26 T=54 |
| Dihyro-progesterone (ng/mL) | High | 0.88 (0.74, 0.95) | 0.47 | 0.78 (0.62, 0.88) | 0.60 | E=11 C=32 T=43 | 0.68 (0.53, 0.79) | 0.69 | 0.57 (0.42, 0.69) | 0.50 | E=24 C=29 T=53 |
| | Low | 0.93 (0.79, 0.98) | | 0.82 (0.66, 0.91) | | E=10 C=35 T=45 | 0.71 (0.57, 0.81) | | 0.64 (0.49, 0.76) | | E=22 C=32 T=54 |
| Epi-Androsterone (ng/mL) | High | 0.89 (0.74, 0.96) | 0.82 | 0.73 (0.55, 0.85) | 0.31 | E=9 C=31 T=40 | 0.64 (0.48, 0.75) | 0.23 | 0.53 (0.38, 0.67) | 0.18 | E=22 C=29 T=51 |
| | Low | 0.91 (0.77, 0.96) | | 0.83 (0.68, 0.92) | | E=11 C=33 T=44 | 0.75 (0.60, 0.84) | | 0.66 (0.52, 0.78) | | E=22 C=30 T=52 |
| Epi-Testosterone (ng/mL) | High | 0.98 (0.84, 1.00) | 0.04 | 0.88 (0.72, 0.96) | 0.07 | E=7 C=34 T=41 | 0.73 (0.59, 0.83) | 0.48 | 0.66 (0.51, 0.78) | 0.34 | E=21 C=32 T=53 |
| | Low | 0.85 (0.70, 0.92) | | 0.73 (0.57, 0.84) | | E=14 C=33 T=47 | 0.66 (0.51, 0.77) | | 0.55 (0.41, 0.68) | | E=25 C=29 T=54 |
| Estrone (ng/mL) | High | 0.88 (0.73, 0.95) | 0.41 | 0.73 (0.56, 0.85) | 0.19 | E=11 C=31 T=42 | 0.66 (0.51, 0.77) | 0.55 | 0.54 (0.39, 0.67) | 0.29 | E=24 C=27 T=51 |
| | Low | 0.93 (0.81, 0.98) | | 0.86 (0.71, 0.93) | | E=10 C=36 T=46 | 0.73 (0.59, 0.82) | | 0.66 (0.52, 0.77) | | E=22 C=34 T=56 |
| Hydroxy Pregnenolone (ng/mL) | High | 0.93 (0.80, 0.98) | 0.42 | 0.83 (0.68, 0.92) | 0.49 | E=10 C=35 T=45 | 0.73 (0.59, 0.83) | 0.40 | 0.62 (0.47, 0.74) | 0.65 | E=23 C=30 T=53 |
| | Low | 0.88 (0.73, 0.95) | | 0.77 (0.60, 0.87) | | E=11 C=32 T=43 | 0.66 (0.51, 0.77) | | 0.59 (0.44, 0.71) | | E=23 C=31 T=54 |
| Non-esterified cholesterol (µg/mL) | High | 0.96 (0.84, 0.99) | 0.08 | 0.86 (0.71, 0.93) | 0.12 | E=11 C=37 T=48 | 0.82 (0.69, 0.90) | <0.01 | 0.72 (0.57, 0.82) | <0.05 | E=20 C=33 T=53 |
| | Low | 0.85 (0.69, 0.93) | | 0.73 (0.55, 0.84) | | E=10 C=30 T=40 | 0.57 (0.42, 0.69) | | 0.50 (0.36, 0.63) | | E=26 C=28 T=54 |
| Pregnenolone (ng/mL) | High | 0.95 (0.82, 0.99) | 0.13 | 0.88 (0.73, 0.95) | 0.08 | E=7 C=37 T=44 | 0.75 (0.61, 0.85) | 0.22 | 0.69 (0.54, 0.80) | 0.12 | E=19 C=34 T=53 |
| | Low | 0.86 (0.71, 0.93) | | 0.72 (0.54, 0.83) | | E=14 C=30 T=44 | 0.64 (0.49, 0.75) | | 0.52 (0.37, 0.65) | | E=27 C=27 T=54 |
| Progesterone (ng/mL) | High | 0.93 (0.81, 0.98) | 0.31 | 0.89 (0.75, 0.95) | <0.05 | E=9 C=38 T=47 | 0.77 (0.63, 0.86) | 0.08 | 0.73 (0.58, 0.83) | <0.05 | E=19 C=34 T=53 |
| | Low | 0.87 (0.72, 0.94) | | 0.69 (0.51, 0.82) | | E=12 C=29 T=41 | 0.62 (0.47, 0.73) | | 0.48 (0.34, 0.61) | | E=27 C=27 T=54 |
| SHBG (nmol/L) | High | 0.89 (0.75, 0.95) | 0.60 | 0.81 (0.66, 0.90) | 0.88 | E=11 C=35 T=46 | 0.77 (0.63, 0.86) | 0.09 | 0.70 (0.56, 0.81) | <0.05 | E=19 C=34 T=53 |
| | Low | 0.93 (0.79, 0.98) | | 0.79 (0.62, 0.89) | | E=10 C=32 T=42 | 0.62 (0.47, 0.73) | | 0.51 (0.37, 0.64) | | E=27 C=27 T=54 |
| T (ng/mL) | High | 0.93 (0.80, 0.98) | 0.44 | 0.85 (0.70, 0.93) | 0.29 | E=10 C=34 T=44 | 0.77 (0.64, 0.86) | 0.11 | 0.71 (0.56, 0.81) | 0.06 | E=20 C=33 T=53 |
| | Low | 0.88 (0.74, 0.95) | | 0.75 (0.59, 0.86) | | E=11 C=33 T=44 | 0.62 (0.47, 0.73) | | 0.51 (0.37, 0.64) | | E=26 C=28 T=54 |

**Supplementary Table 15 Demographics in TMA**

| | | AA | EA | Overall | p-value |
|---|---|---|---|---|---|
| Overall | N | 107 (44.6) | 133 (55.4) | 253 (100%) | |
| Age at Diagnosis | Mean/Std/N | 56.0/7.1/107 | 59.7/6.7/133 | 58.1/7.1/240 | <.001 |
| | Median/Min/Max | 55.0/41.0/70.0 | 60.0/41.0/78.0 | 58.0/41.0/78.0 | |
| Age* | < 55 | 46 (43.0%) | 29 (21.8%) | 80 (31.6%) | <.001 |
| | >= 55 | 61 (57.0%) | 104 (78.2%) | 173 (68.4%) | |
| Tobacco Use | None | 42 (39.3%) | 49 (37.4%) | 92 (38.5%) | <0.01 |
| | Former | 32 (29.9%) | 63 (48.1%) | 95 (39.7%) | |
| | Current | 33 (30.8%) | 19 (14.5%) | 52 (21.8%) | |
| pre-RP PSA | Mean/Std/N | 9.1/10.3/106 | 9.1/10.7/132 | 9.1/10.5/238 | 0.48 |
| | Median/Min/Max | 6.5/0.7/82.1 | 6.3/0.1/87.9 | 6.3/0.1/87.9 | |
| PSA* | < 4 | 16 (15.1%) | 24 (18.2%) | 40 (16.7%) | 0.92 |
| | 4 to <10 | 70 (66.0%) | 82 (62.1%) | 152 (63.6%) | |
| | 10 to <20 | 12 (11.3%) | 16 (12.1%) | 28 (11.7%) | |
| | >= 20 | 8 (7.5%) | 10 (7.6%) | 19 (7.9%) | |
| Grade | Grade II | 32 (29.9%) | 33 (24.8%) | 65 (25.7%) | 0.39 |
| | Grade III | 75 (70.1%) | 100 (75.2%) | 188 (74.3%) | |
| Clinical Gleason | <= 3+4 | 61 (57.0%) | 76 (58.0%) | 138 (57.7%) | 0.90 |
| | >= 4+3 | 46 (43.0%) | 55 (42.0%) | 101 (42.3%) | |
| Clinical T-stage | cT1 | 71 (68.9%) | 90 (68.7%) | 162 (68.9%) | 0.53 |
| | cT2 | 28 (27.2%) | 39 (29.8%) | 67 (28.5%) | |
| | cT3 | 4 (3.9%) | 2 (1.5%) | 6 (2.6%) | |
| Path Gleason | < 3+4 | 45 (42.5%) | 49 (37.1%) | 94 (39.3%) | 0.43 |
| | > 4+3 | 61 (57.5%) | 83 (62.9%) | 145 (60.7%) | |
| Path T-stage | pT2 | 71 (68.9%) | 75 (57.7%) | 146 (62.4%) | 0.23 |
| | pT3 | 27 (26.2%) | 46 (35.4%) | 74 (31.6%) | |
| | pT4 | 5 (4.9%) | 9 (6.9%) | 14 (6.0%) | |
| Persistent Disease? | No | 84 (78.5%) | 106 (79.7%) | 200 (79.1%) | 0.87 |
| | Yes | 23 (21.5%) | 27 (20.3%) | 53 (20.9%) | |
